# An Examination of an Enhanced Remote Sensing Method for Agent Attribution of Forest Disturbance

**DOI:** 10.1101/2020.11.23.394221

**Authors:** Hugh Marshall Worsham

## Abstract

Patterns of disturbance in Sierra Nevada forests are shifting as a result of changing climate and land uses. These changes have underscored the need for a monitoring system that both detects disturbances and attributes them to different agents. Addressing this need will aid forest management and conservation decision-making, potentially enhancing forests’ resilience to changing climatic conditions. In addition, it will advance understanding of the patterns, drivers, and consequences of forest disturbance in space and time. This study proposed and evaluated an enhanced method for disturbance agent attribution. Specifically, it tested the extent to which textural information could improve the performance of an ensemble learning method in predicting the agents of disturbance from remote sensing observations. Random Forest (RF) models were developed to attribute disturbance to three primary agents (fire, harvest, and drought) in Stanislaus National Forest, California, U.S.A., between 1999 and 2015. To account for spectral behavior and topographical characteristics that regulate vegetation and disturbance dynamics, the models were trained on predictors derived from both the Landsat record and from a digital elevation model. The predictors included measurements of spectral change acquired through temporal segmentation of Landsat data; measurements of patch geometry; and a series of landscape texture metrics. The texture metrics were generated using the Grey-Level Co-Occurrence Matrix (GLCM). Two models were produced: one with GLCM texture metrics and one without. The per-class and overall accuracies of each model were evaluated with out-of-bag (OOB) observations and compared statistically to quantify the contribution of texture metrics to classification skill. Overall OOB accuracy was 72.0% for the texture-free model and 72.2% for the texture-dependent model, with no significant accuracy difference between them. Spatial patterns in prediction maps cohered with expectations, with most harvest concentrated in mid-elevation forests and fire and stress co-occurring at lower elevations. Altogether, the method yielded adequate identification of disturbance and moderate attribution accuracy for multiple disturbance agents. While textures did not contribute meaningfully to model skill, the study offers a strong foundation for future development, which should focus on improving the efficacy of the model and generalizing it for systems beyond the Central Sierra Nevada.

## 1. Introduction

Disturbance regulates the composition and structure of temperate forests by altering processes of vegetation growth, death, decomposition, and regeneration (Turner 2010). Disturbance agents interact with pre-disturbance conditions to produce variable effects with profound consequences for post-disturbance regeneration (Collins and Roller 2013, Coop et al. 2016, Shive et al. 2018), as well as for carbon storage, water cycling, timber productivity, wildlife habitat, and other ecological goods and services that forests provide.

Consider three examples. In the highest-intensity regions of a wildfire, living trees of all age classes may be carbonized or left as standing or downed deadwood, while organic material is consumed from the surface through much of the root zone (Cochrane and Ryan 2009, Perry et al. 2011). On the lower-intensity margins, fire may thin the understory or selectively kill weakened individuals and more vulnerable species, freeing resources that enable mid- to late-seral species to release (Braziunas et al. 2018). A clear-cut harvest, in turn, abruptly removes most or all tree cover, leaving few or no standing stems (Franklin et al. 2002, Tappeiner et al. 2015). Post-harvest regeneration must begin from “the ground up,” via an existing seedbank or artificial seeding or planting. On the other hand, some forest disturbances are less abrupt. Mortality due to desiccation stress or beetle infestation typically unfolds over months or years, often with species or age-class selectivity. Infestations yield relatively slow declines in chlorophyll canopy content, frequently yielding distinctive “red” and “gray” phases of decline, and standing dead stems may remain on site for many years (Ciesla 2000). When salvage logging is not applied, much of the nutrient stock may also be retained on site, as dead stems fall and decompose, but the site may face an increased wildfire risk (Tappeiner et al. 2015, Larvie et al. 2018).

From a theoretical perspective, what constitutes a disturbance, and how disturbances ought to be differentiated from other kinds or degrees of perturbation that affect biological communities, have proven thorny questions to answer (Sousa 1984). Disturbances are often labeled “natural” (e.g., fire, windthrow, flood, pest infestation, drought) or “anthropogenic” (e.g., biological invasion, forest management treatment, fragmentation, roadcut, plantation-conversion) (Dale et al. 2001, Turner and Gardner 2015). However, insisting on a sharp line between these categories is unrealistic. In the western United States, as in other parts of the world, the legacies of indigenous landscape management practices likely cannot be disentangled from the region’s “natural” fire regime (Conedera et al. 2009, Trauernicht et al. 2015). Equally, anthropogenic climate change appears to be influencing “natural” disturbance processes such as desiccation stress and dieback in western U.S. forests (Clark et al. 2016).

Two further problems beset theoretical characterizations of disturbance. First, without some qualification, the concept implies a possibility of stasis that rarely occurs in natural systems (Connell and Sousa 1983, Sousa 1984). Many biological communities readily shift among a set of alternative stable states (Beisner et al. 2003), or even alternative transient states (Fukami and Nakajima 2011). In the absence of an objective way to identify where a system lies within its alternative-state frontier at any moment, it is hard to say when a perturbation is disruptive enough to qualify as a disturbance. Second, disturbance agents often interact: to cite one example, drought stress can inhibit trees’ defenses against infections and parasites, in addition to rendering them more vulnerable to fire. (Anderegg et al. 2015, Johnstone et al. 2016, Seidl and Rammer 2017, Simler et al. 2018). To attempt to differentiate particular agents as proximate or ultimate causes of disturbance often seems more a hermeneutic exercise than an empirical one.

Considering these difficulties, while acknowledging that disturbance is nevertheless a useful way to describe a class of environmental phenomenon, this paper holds with the idea that disturbance “lies near one extreme of the continuum of perturbations that affect organisms” (Sousa 1984). It also assumes, *sec.* Peters et al. (2011), that an adequate description of a given disturbance needs to account for at least: (1) the environmental agent(s) of disturbance (“drivers”), (2) structural and functional characteristics of the system prior to the disturbance (“initial system properties”), and (3) interactions between the first two components that give rise to physical, chemical, and biological mechanisms of change (“mechanisms”). An adequate description of disturbance should also consider the consequences explicitly. The outcomes of forest disturbance can include vegetation morbidity and mortality in the short-term and changes in age class, species composition and dominance, hydrologic function, or ecosystem state (among others) in the long-term. For this paper’s purposes, I take forest disturbance to mean a discrete application of energy to, or expenditure of energy within, a forested landscape that results in mortality, morbidity, or displacement of vegetation and that opens opportunities for the establishment of new individuals. I am primarily concerned with disturbances observable at the hectare scale (10,000 m^2^) and larger, because of size of the area of study (∼3600 km^2^) and the spatial resolution of the observations used (∼900 m^2^).

### 1.1. Forest disturbance and climate change

A growing body of evidence suggests that patterns of disturbance in the forests of the Sierra Nevada of California are shifting (Breshears et al. 2005, Millar et al. 2007, Adams et al. 2010, Allen et al. 2010, Cohen et al. 2016). For instance, timber harvesting in national forests has decreased since the 1970s, while the incidence of wildfire and pest infestation has increased (Oswalt et al. 2019). In the Sierra Nevada, desiccation stress was widespread during the 2012–2015 drought, but it was also attended in some areas, such as Sequoia & Kings Canyon National Parks, by severe outbreaks of western and mountain pine beetles (Larvie et al. 2018).

So far, one consistent net effect of these shifts is high tree mortality (Potter 2017, Crockett and Westerling 2018, Fettig et al. 2019). The new dynamics may also be inducing species shifts and biodiversity losses (Paz-Kagan et al. 2017), and they could drive the replacement of forests by non-forest land cover types, such as shrubland or meadow (Thorne et al. 2017, 2018). Some have projected that these trends will continue as a result of climate change and anthropogenic activity, with consequent impacts on the services that currently forested landscapes provide.

Given these trends, advancing understanding of the patterns, drivers, and consequences of forest disturbance in space and time is a research priority (Trumbore et al. 2015, Johnstone et al. 2016). McDowell et al. (2015) note a “lack of a comprehensive monitoring system” that can both identify terrestrial disturbances and attribute them to specific agents. Filling this gap would help forest resource managers understand how forests respond to changing climatic conditions. In addition, reliable quantification of historical and emerging disturbances will help to improve the skill of empirical models of spatial pattern, population dynamics, forest regeneration, carbon storage, and water cycling. In the long run, this effort could also improve the prospects for quantitative description of ecological disturbance in the context of alternative stable (or transient) states by improving the resolution of pre- and post-disturbance baselines. Finally, it will support strategies for conserving, restoring, or adaptively transitioning forests in areas facing increasing vulnerabilities to various agents of disturbance (Millar et al. 2007, Hansen and Turner 2019).

### 1.2. Review of remote sensing methods for forest disturbance detection and attribution

Over the past decade, efforts to detect and attribute forest change have proliferated. However, the field has yet to settle on a set of approaches that produce reliable estimates that can be compared across disturbance regimes or regions. The field currently comprises a somewhat incongruent set of algorithms, satellite and aerial monitoring platforms, and field assessment protocols. The USDA Forest Service’s Forest Inventory and Analysis (FIA) program provides data on most classes of forest-disturbance agent, but only across a network of sampling plots (Schroeder et al. 2014). In California, the dataset extends back to 2001, with repeat surveys conducted on each plot approximately once a decade (Christensen et al. 2016). This relative infrequency, along with the FIA’s policy of obscuring the precise locations of most sample plots, substantially limits the data’s suitability for quantifying spatially continuous change. Aerial insect and disease detection surveys are similarly discontinuous and coarsely resolved in space and time. In response to these limitations, researchers and managers have increasingly turned to satellite remote sensing methods for their ability to capture a wide range of spatial and temporal variability across large regions.

The history of remote sensing methods for forest change detection extends at least as far as the 1920s, when an entomology study analyzed oblique aerial photography to identify spruce budworm mortality in Canadian spruce forests (Ciesla 2000). This process was improved substantially by a double-sampling approach developed in the 1950s and 1960s, in which tree mortality estimates were made through stereoscopic interpretation of a large sample of photographs and scaled up through statistical comparison with a smaller sample of ground plots (Heller et al. 1959, Wear et al. 1966). Double sampling allowed for statistically valid estimations of canopy loss across wide geographic areas (Lund 1997). With the increasing availability of color and infrared film, researchers and forest managers also began to exploit spectral information to identify crown fade and red and grey phases of beetle infestations (Hadfield 1968, Hanson and Lautz 1971). With the launch of the first civilian Earth-observing satellites, Landsat I in 1972, Geostationary Operational Environmental Satellite (GOES-1) in 1975, and the Advanced Very High Resolution Radiometer (AVHRR) in 1978, remote sensing methods for forest change detection boomed.

Methods developed early on—and still in widespread use today—derive information by comparing two or more images made at separate points in time over the same geographic area. In pre-classification change detection methods, analysts compare the images’ raw spectral data. In post-classification techniques, each pixel in an image is assigned to a defined land-cover type, and intertemporal differences are evaluated as changes in class (Iverson et al. 1989). While post-classification approaches allow for the integration of multiple data types and can minimize the effects of exogenous atmospheric or radiometric distortions, they can carry an additional error burden due to information loss in the classification procedure (Coops et al. 2007). More complex approaches in this category involve principal component analysis, in which correlated spectral returns are compressed so that change detection is performed on independent linear transformations of the original data, and change vector analysis, which decomposes spectral responses into magnitude and directional components (Fung and Ledrew 1987, Lu et al. 2004, Khorram et al. 2016). In addition to uncovering dramatic vegetation changes around the world, these two-date change-detection approaches identified two important requirements for any attempt at forest change detection. First, special care must be taken to align the images geometrically and radiometrically as nearly as possible to avoid false-negative change detection as a result of registration inconsistencies. Second, it is imperative to account for seasonal change, either through multi-band analysis, index computation, or seasonal compositing of multiple images, as phenological change can easily be confused with stress-related change, particularly in visible wavelengths (Khorram et al. 2016).

Because vegetation disturbance is a dynamic process operating on multiple timescales, two-date comparison methods carry obvious limitations. In the mid-2000s, a suite of algorithms was developed to address this gap; these operate on the unique time signatures that different directions and magnitudes of vegetation change leave behind (García-Haro et al. 2001, Potter et al. 2007, Goodwin et al. 2008, Vogelmann et al. 2009, 2012). Such approaches are able to detect either abrupt changes (anomalies) or longer-duration changes (trends) with suitable accuracy for monitoring and management. The most sophisticated of these is the Vegetation Change Tracker (VCT) (Huang et al. 2010). This method characterizes the temporal profile of each pixel in a time series stack and classifies the pixels into one of several types (persistent forest, persistent non-forest, disturbed forest, or regenerating forest), based on comparisons of absolute change in the series with predetermined thresholds. Although these improve on two-date methods, as Kennedy et al. (2010) point out, anomaly-targeted algorithms tend to exclude long-term trend changes as noise, while trend-targeted algorithms do the same for abrupt anomalies.

Since 2010, a new generation of algorithms has emerged to disentangle remotely sensed time series data to capture both abrupt disturbance events and longer-phase trend change. These temporal segmentation algorithms are summarized in Table 1.

**Table 1.**
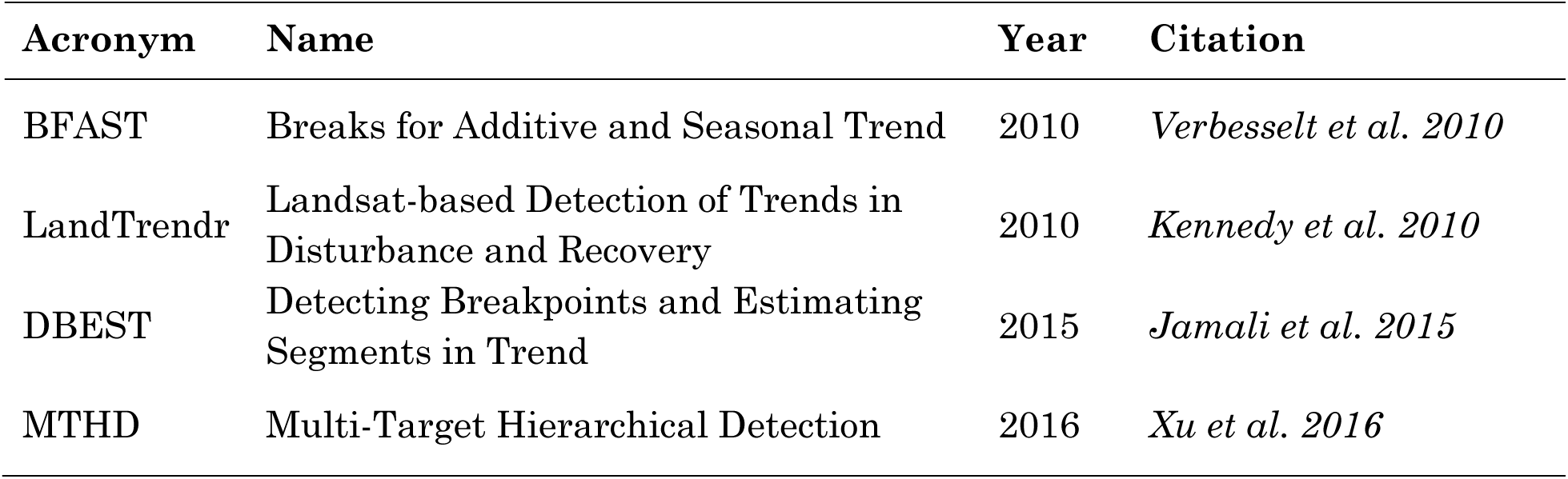
Recent temporal segmentation procedures for discriminating abrupt and trend vegetation change using remotely sensed data.

| Acronym | Name | Year | Citation |
| --- | --- | --- | --- |
| BFAST | Breaks for Additive and Seasonal Trend | 2010 | *Verbesselt et al. 2010* |
| LandTrendr | Landsat-based Detection of Trends in Disturbance and Recovery | 2010 | *Kennedy et al. 2010* |
| DBEST | Detecting Breakpoints and Estimating Segments in Trend | 2015 | *Jamali et al. 2015* |
| MTHD | Multi-Target Hierarchical Detection | 2016 | *Xu et al. 2016* |

Although the methods differ somewhat in their implementation, in general they apply a statistical model, such as linear segmentation, Fourier curve fitting, or quadratic smoothing, to a time-series stack of remotely sensed data in order to derive information about each pixel’s spectral trajectory over time. The output is usually an array that includes a per-pixel estimate of the location, timing, duration, and in some cases, magnitude of spectral change. Based on these outputs, an analyst can inquire about patterns of behavior among pixels or among aggregates formed based on the adjacency or similarity of pixels in one or more dimensions.

In the U.S., researchers and managers have deployed both two-date image differencing and the more complex algorithmic approaches in several prominent disturbance monitoring programs. Monitoring Trends in Burn Severity (MTBS) uses a two-date procedure on Normalized Burn Ratio (NBR) values from Landsat images to map the severity of large fires (Eidenshink et al. 2007). The ForWarn system uses Normalized Difference Vegetation Index (NDVI) anomaly calculated on MODIS data to model disturbances in near-real-time. However, its coarse spatial resolution (minimum 250 m) makes it insensitive to finer-scale disturbances, including most management treatments on public forested lands in the western U.S. (Hargrove et al. 2009). The LANDFIRE disturbance database resolves 12 disturbance agent classes and dozens of sub-classes at 30- m spatial resolution (Rollins 2009, Vogelmann et al. 2011). The program draws on multiple algorithmic, remote sensing, and *in situ* data sources, including MTBS and VCT, but its evidently incomplete record dates back only to 1999. The North American Forest Dynamics program has leveraged the VCT algorithm to build a wall-to-wall map of U.S. forest disturbance across the entire Landsat record at 30-m scale (Goward et al. 2016).

The efforts above have advanced *detection* of forest disturbance; methods for *agent attribution*, on the other hand, remain embryonic. The most reliable approaches require extensive technician analysis of multiple datastreams, including *in situ* observations, management treatment records, and aerial- and satellite-platform sensing (e.g., Schmidt 2014, Cohen et al. 2016). The process is time-intensive and beset with error when multiple forest types are under investigation, when multiple disturbance agents are active, and when a site experiences more than one disturbance in the same period of analysis. Automating this process through empirical modeling may help to reduce time and resource requirements, in addition to improving accuracy of retrospective analyses and enhancing the relevance of near-real-time disturbance detection and monitoring.

To date, a limited number of efforts at further automating agent attribution have been published. The methods in this paper are heavily indebted to these projects. Neigh et al. (2014) used multiple indices derived from AVHRR and Landsat products in a decision-tree classification to map insect kill and harvest in northern forests of Wisconsin and Minnesota. They achieved per-class accuracy of 65–70% and overall accuracy of 72%. However, the same method applied to forests in the Pacific Northwest yielded inferior results, indicating a need for further refinement before generalizing across forest types. Kennedy et al. (2015) combined LandTrendr temporal segmentation of the 1984–2014 Landsat stack with a Random Forest classification approach (Breiman 2001) to map multiple agent classes in the Pacific Northwest. Oeser and colleagues (2017) applied a similar approach in Central European temperate forests, applying BFAST temporal segmentation to identify abrupt forest loss and passing the resulting spatio-temporal information into a Random Forest classification. They identified harvest, windthrow, cleared windthrow, and bark beetles to 76–86% accuracy. Schroeder et al. (2017) classified fire, harvest, conversion, wind, and drought stress with a Random Forest model trained on VCT and ancillary geophysical variables. Their approach yielded high accuracy (69–86%) across ten Landsat scenes made over various ecoregions of the United States. Interestingly, accuracy was higher when information about the timing (year) of disturbance was excluded from the model. Shimizu et al. (2017) also used Random Forest to classify patches of contemporaneously disturbed pixels to discriminate anthropogenic forest changes, such as logging, plantation conversion, and urbanization in Myanmar. Finally, Shimizu et al. (2019) evaluated the relative effectiveness of several different approaches to disturbance-agent classification in a South Asian tropical forest: threshold-based detection using one spectral index, machine-learning methods trained on temporally segmented vegetation index values, and one machine-learning method trained directly on the Landsat time series without prior temporal segmentation. They found that direct prediction performed better than approaches that included temporal segmentation, with considerable savings in complexity and computational expenditure, but it remains to be seen whether this approach works as well as two-stage methods in other forest types.

Clearly, there is enthusiasm for solving the agent-attribution problem, and the early work suggests that remotely sensed information shows promise for resolving different types of forest disturbance into distinct classes. But much remains to be done to achieve a generalizable method. Some key areas for development include:

1. identifying the most effective combination of spectral bands and indices for accurate modeling across landscape types;
2. determining whether a two-stage method (temporal segmentation plus classification) or a one-stage method (direct classification without temporal segmentation) consistently yields higher accuracy;
3. making use of new spectral measurements of solar-induced chlorophyll fluorescence (SIF), such as the NASA Orbiting Carbon Observatories’ (OCO-2 and OCO-3) SIF products and the Near-Infrared Reflectance of Vegetation (NIRv) index; and
4. assessing whether textural information derived from Landsat scenes can improve classification results.

Here, I address the last two of these prompts by making novel use of NIR_V_ and by testing the contribution of textural information to agent-attribution accuracy.

### 1.3. NIR_V_

This study marks the first use of NIR_V_ in a change-detection procedure. NIR_V_ directly measures the fraction of near-infrared reflectance attributable to chlorophyll, yielding accurate estimates of photosynthesis rate and gross primary production (GPP) (Badgley et al. 2017, 2019, Wu et al. 2020). NIR_V_ tends to be more sensitive to decreases in photosynthetic capacity than other vegetation indices, which offers reason to think that a model trained on NIR_V_ may improve detection of sublethal phases of stress-related disturbances (e.g., drought, beetle infestation).

### 1.4. Texture

On texture, as this paper’s introduction describes, different agents and intensities of disturbance leave different structural legacies on forested landscapes. Vegetation structure, in turn, has been shown to resolve well in textural patterns derived from optical remote sensing measurements (Wood et al. 2012, Lam et al. 2013). At the most basic level, *texture* describes certain spatial properties of a surface—in ordinary experience, these include properties such as smoothness, coarseness, or sharpness. Quantifying texture for analytic purposes is a matter of measuring and expressing differences between high and low points on a surface (*z* differences in Cartesian space), and how near or far those points are from one another (*x-y* differences). Smoother surfaces tend to have smaller *x-y-z* differences, while rougher surfaces tend to have larger differences. Smoothness and coarseness are only two of many relevant textural properties that can be measured statistically from images of a surface. One widely deployed set of metrics is that derived from the Gray Level Co-occurrence Matrix (GLCM) (Haralick et al. 1973). The GLCM procedure tabulates the frequency of co-occurrence of pixel brightness values in adjacent pixels using a set of moving-window comparisons. These frequencies are then used to compute a set of 14+ distinct measurements of texture.

At the pixel level, GLCM metrics describe second-order statistical properties. First-order information, such as the spectral reflectance intercepted by a sensor and recorded as pixel values, generally measures physical behavior (reflectance of electromagnetic energy) or chemical activity (photosynthesis) or a statistically verifiable proxy for the same. GLCM, on the other hand, quantifies relationships between *pairs* of pixels (Hall-Beyer 2017). First- and second-order metrics tend to be statistically independent of each other and so can contribute complementary information to landscape analyses. GLCM textures have long been applied in combination with other variables to improve accuracy of land-use and land-cover classification (Coburn and Roberts 2004). The applicability of GLCM metrics for forest structure analysis has been demonstrated using both high-resolution (IKONOS) and moderate-resolution (SPOT, Sentinel-1, ASTER, Landsat) data, though it likely has much lower utility in land cover applications at spatial resolutions lower than ∼50 m per pixel (Marceau et al. 1990, Ozdemir et al. 2012, Wood et al. 2012).

Textural metrics have the potential to improve disturbance agent attribution because of the relationships between agent and stand structure on the ground, and between stand structure and texture in images. These relationships may be especially important in the Central Sierra Nevada forests evaluated in this study because of the agents that are most prevalent in the region: fire, drought, insects, and harvest. These tend to leave visually distinctive and analytically differentiable patterns on the landscape (Fig. 1) and may contribute decision-enhancing information to an agent-attribution modeling procedure.

**Figure 1.**
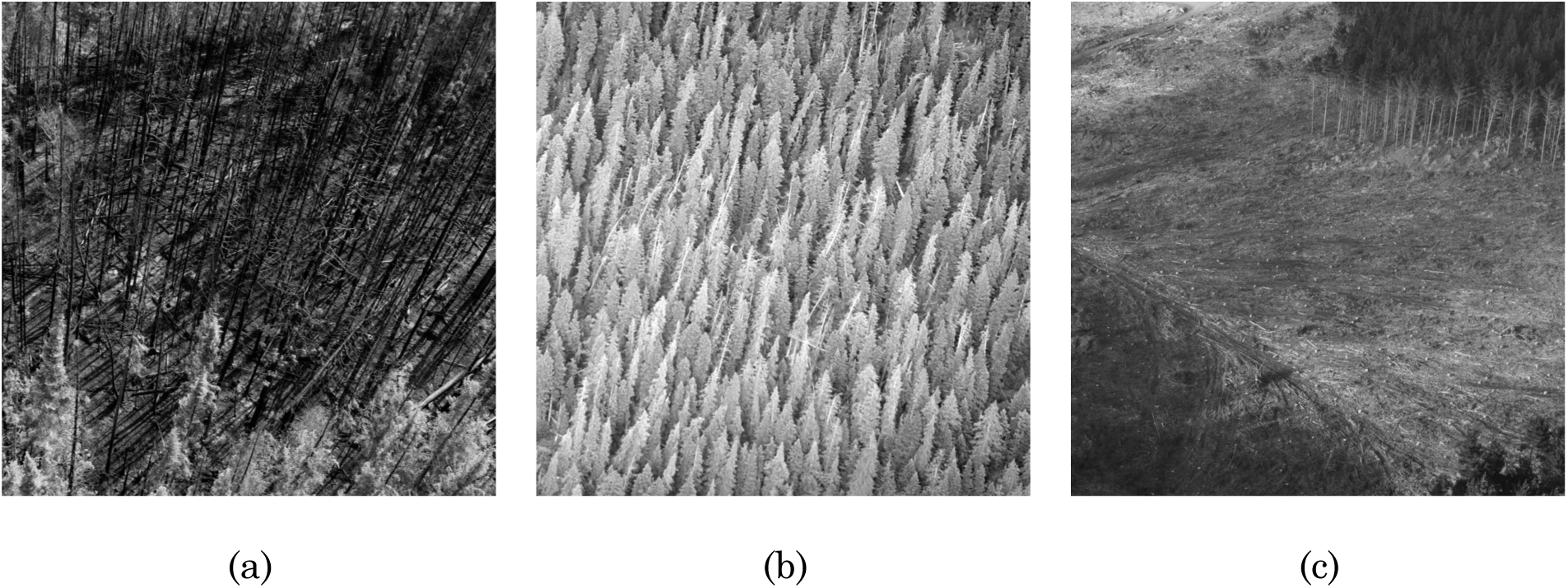
Photographs of Sierra Nevada mixed-conifer forest sites disturbed by (a) mixed-severity fire, (b) bark beetles, and (c) harvest. Each photograph was made within one year of disturbance and reveals a distinctive structural legacy.

### 1.5. Motivation

California’s Central Sierra Nevada offers valuable opportunities to study forest disturbance and its drivers using the emerging remote sensing methods described above. Observed climatic changes, including warmer winter and spring temperatures, precipitation shifts from snow to rain, lower peak snowpack depth, and early spring drydown, have been documented across the region (Vicuna and Dracup 2007). These shifts are connected to a multitude of forest changes. For example, whitebark pine (*Pinus albicaulis* Engelm.) and ponderosa pine (*Pinus ponderosa*) have experienced widespread mortality due to mountain pine beetle, western pine beetle, and desiccation stress (Millar et al. 2012, Birdsey et al. 2019). The region has also faced compositional shifts and increases in stem density in mid-elevation coniferous stands, as well as canyon oak regeneration in stands previously occupied by conifers (Dolanc et al. 2013, 2014).

The site selected for study, Stanislaus National Forest, is an archetype of these trends. Fire, harvest, thinning, drought, and insect stress have been extensive and well distributed across elevational gradients over the past three decades. The prevalence of disturbance makes it a prime site for an attempt at complex disturbance agent attribution. Indeed, the Forest has been a subject of at least two prior disturbance-detection studies using LandTrendr and VCT, respectively (Schmidt 2014, Birdsey et al. 2019). Both relied on manual interpretation of multiple data sources for agent attribution. Schroeder et al.’s (2017) semi-automated approach using VCT and Random Forest classification included one Landsat tile that partially overlapped the Forest. Their results showed agent classification agreement above 90 percent for the Sierra Nevada site, indicating strong potential for this approach in the region.

The overarching aim of this project was to test whether a Random Forest ensemble learning method for classifying forest disturbance agents at the 30-m Landsat pixel scale can be improved by incorporating textural information.

### 1.6. Research questions

1. To what extent does the inclusion of textural information improve attribution of the agents of disturbance in Stanislaus National Forest?
2. What are the relative contributions of three independent textural metrics to classification accuracy?

### 1.7. Study objectives

1. <u>Agent-attribution model:</u> Evaluate the capacity of an ensemble learning method to classify Landsat-derived pixel data according to three agent-based forest disturbance classes (fire, harvest, stress) and stable forest/non-forest.
2. <u>Texture contribution to accuracy:</u> Assess the per-class and overall accuracy of texture-free and texture-dependent agent models to evaluate whether a model with textural metrics is more effective at identifying agents of disturbance than one without.
3. <u>Variable importance:</u> Determine which predictor variables are most useful for attributing forest disturbance agents in a Sierra Nevada forest.

## 2. Methods

### 2.1. Study site

Stanislaus National Forest is a 3,634-km^2^ federal landscape administered by the USDA Forest Service on the western slope of the Sierra Nevada in California (Fig. 2). The forest abuts the northern border of Yosemite National Park and contains three federal Wilderness areas (Mokelumne, Carson-Iceberg, and Emigrant) to the north and east. The region’s climate is Mediterranean, with average precipitation around 125 cm (990 cm equivalent snowfall.) The jurisdiction spans a broad elevational gradient, from 450 m in the western foothills to over 3350 m near the Sierra crest. The Forest contains more than 1200 km of rivers and streams. Vegetative communities include oak woodlands at lower elevations, mixed conifer forests at middle elevations, and subalpine vegetative communities at the highest elevations.

**Figure 2.**
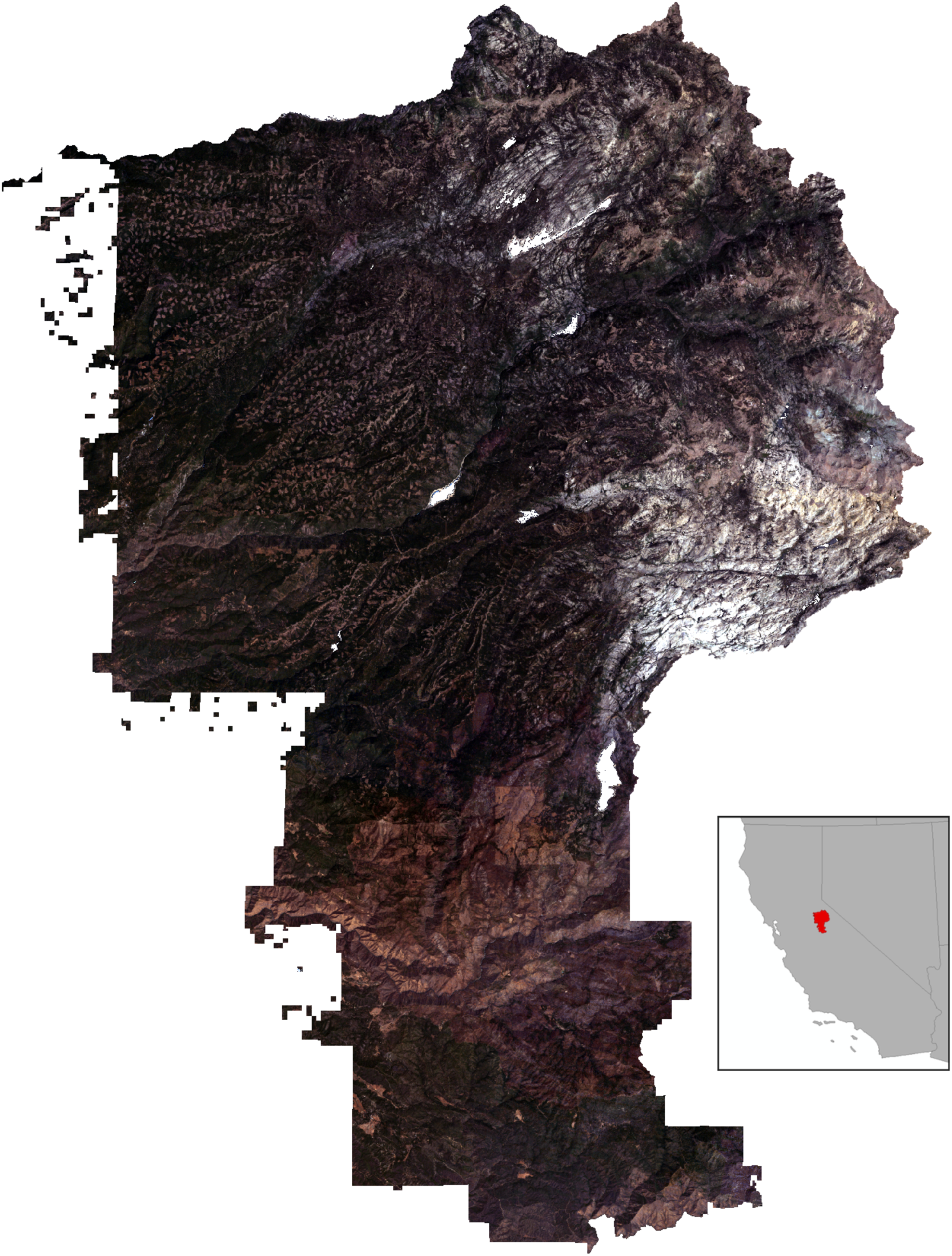
True-color composite image of Stanislaus National Forest in July 2014 (California, U.S.A., inset). The composite was created from bands 2–4 of a Landsat 8 Enhanced Thematic Mapper (ETM+) image made approximately eleven months after the Rim Fire began. The fire scar is visible across the image’s lower third. Extensive harvest patches (∼16 hectares each) appear in the speckled regions to the north and west. Surface water and cloud shadows are masked and appear white.

Since 2000, the Forest has experienced two major wildfires: the 2013–2014 Rim Fire, which burned 257,314 acres and the 2018 Donnell Fire, which burned 36,450 acres. Forest ecosystems in the domain are subject to other natural disturbance regimes, such as conifer beetle eruptions, severe winter wind events, and avalanches. They are also harvested for merchantable timber and thinned for fire resistance, pest management, species selection, and site productivity; these operations often register as vegetation loss in change-detection analyses, but because of forest management practice guidelines, are typically constrained to clearly delineated areas less than 16 hectares (0.16 km^2^).

### 2.2. Data preparation

#### 2.2.1. Reference data

Ideally, reference data for model training and accuracy assessment would come from data acquired in the field. However, a consistent, spatially explicit longitudinal record of *in situ* disturbance observations does not exist for California, and due to time and cost constraints I was unable to assemble such a record myself. Instead, I used the Landscape Fire and Resource Management Planning Tools (LANDFIRE) Disturbance Public Model-Ready Events Geodatabase. In its original form, this dataset comprises a set of polygon shapefiles demarking the locations, extents, types, and timing of disturbances and management treatments. The polygons are submitted annually to LANDFIRE, a joint program of the USDA Forest Service and U.S. Department of the Interior, by contributors from federal and state resource management agencies, private organizations, and national/regional fire mapping programs, such as MTBS and CalFire’s Fire and Resource Assessment Program (FRAP). Data submissions must meet minimum standards for inclusion, and they are subsequently analyzed for positional accuracy and quality and then corrected for topological inconsistencies. In the Model Ready Events dataset, the set of polygons is reduced to portray one unique event per location per year between 1999 and 2014, using a hierarchical decision procedure. The polygons that comprise the final dataset are those with the greatest-magnitude impact on vegetation.

Because these data were used for training as well as validation, the model inherits error from the reference set. However, LANDFIRE currently offers the most extensive and longest record of disturbances and treatments available for the study area. (The dataset also offers nearly full coverage of the continental United States, which would aid testing of the generalizability of the methods in this study in the future). Moreover, with the exception of data generated by MTBS, the records are created without reliance on Landsat observations. In the study area, because of the extensive records maintained by CalFire FRAP, none of the fire event polygons were derived from MTBS. It stands to reason that the reference set is as independent of the predictor data as is feasible. I considered the reference set sufficient in light of the fact that this is foremost a proof-of-concept study but acknowledge that more reliable training and reference data would improve the reliability of the model.

I converted the polygons to Geotiff raster format and randomly selected 200 sample points from each of five strata: fire, harvest/treatment, stress, stable forest, and stable non-forest. For each of the disturbed points, I preserved the year reported and the assigned disturbance agent from LANDFIRE. In some cases, the sampling yielded multiple disturbances per point during the time interval. When this happened, I selected the highest-severity disturbance in the set to maintain consistency with the procedure in the temporal-segmentation algorithm described in *§2.2.2* below. The 1000 total sample points represented three classes of disturbance agent (fire, harvest, stress) and two classes of stability (stable forest, stable non-forest).

#### 2.2.2. Landsat Tier-1 surface reflectance datasets

A flowchart of the remaining analytical steps appears in Fig. 3. The next task was to generate predictor variables for training the classification model. Using the Google Earth Engine API (Gorelick et al. 2017), I assembled an image collection of Landsat 5 Thematic Mapper (TM), Landsat 7 Enhanced Thematic Mapper (ETM+) and Landsat 8 Operational Land Imager (OLI) datasets acquired over the study area, preprocessed to Tier 1 surface reflectance. ETM+ and OLI data provide moderate-to-high spatial and temporal resolution (30 meters per pixel for non-thermal bands on a 16-day return interval). They offer adequate spectral resolution for vegetation change detection.

**Figure 3.**
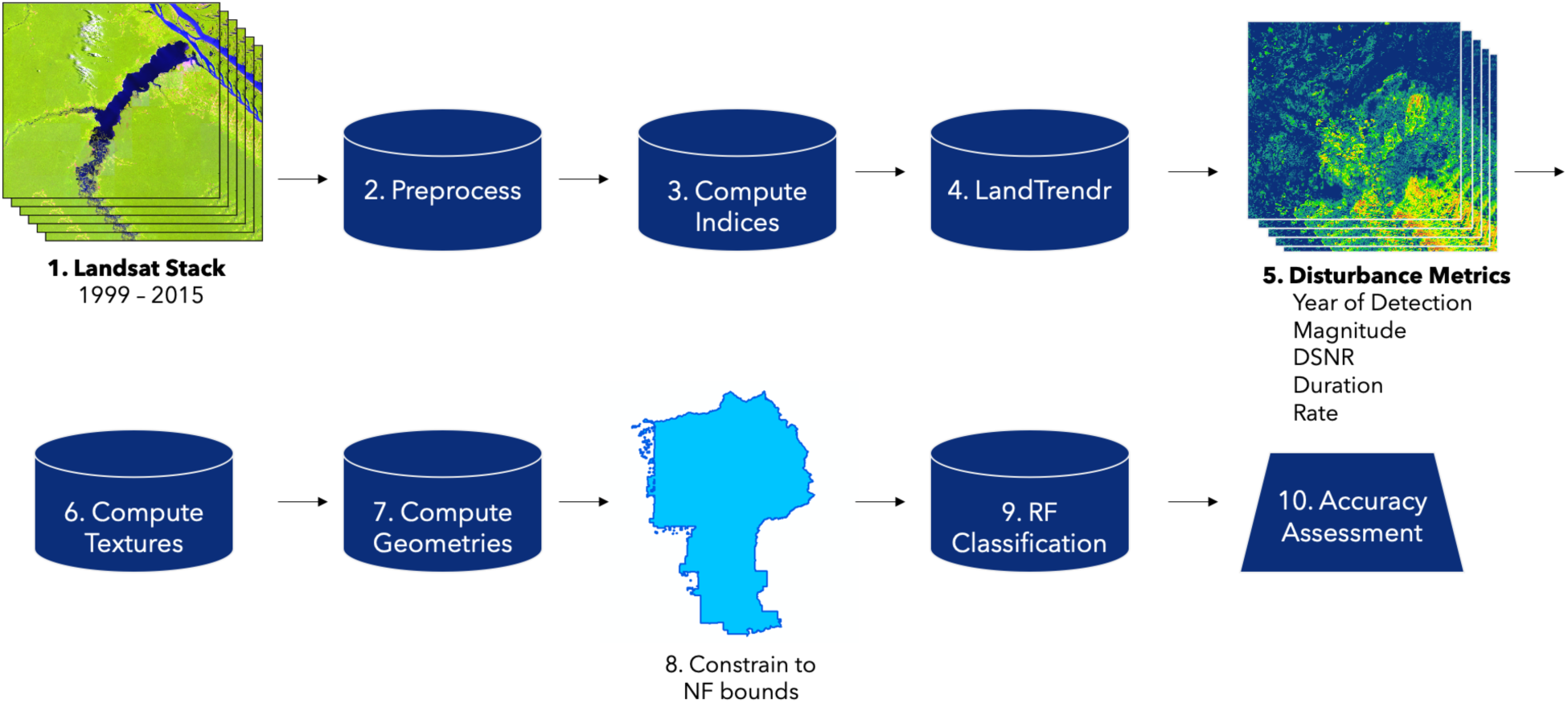
A flowchart of the data processing methods detailed in this study. Steps 1–3 pertain to §2.2.2. Step 4 is described in §2.2.3. Steps 5–8 are detailed in §2.2.4 and §2.2.5. Step 9 is described in §2.3, and Step 10 in §2.4.

The image collection included all images made during the peak growing season (June 21 – September 20) in the years between 1999 and 2015, inclusive. Next, to standardize the ETM+ and OLI data, I applied a slope-intercept harmonization algorithm, which normalized OLI surface reflectance values to ETM+ values. A detailed discussion of this procedure is available in Roy et al. (2016). I then applied a masking function to each image using the pixel-QA band to remove clouds, snow, cloud shadows, and water, in order to avoid generating outlier band ratio values that could lead to false positive change detection. The final collection contained 373 images in total, between nine and 31 per year for an average of 22 images per year.

The next step was to build a summary dataset of annual surface reflectance images. I computed the medoid value per pixel from the annual subsets of masked image collections. The medoid is a measure of center that minimizes the vector distance to all other points in the set. In an odd set in one-dimensional number space, this is the median. In an even set, in which the median would fall in the interval between two values, the medoid is constrained to one of the values actually present in the dataset. In this case, the medoid computation selected the lower value in the interval. Medoid compositing produced 17 images, one for each year in the study period.

The consequence of this compositing was that subsequent analysis evaluated interannual change, a significant scaling up from the 16-day temporal resolution of the original Landsat data. There are tradeoffs in any data selection procedure. While some information is compressed or lost in generating annual medoids, this process reduces the error in intertemporal comparison resulting from radiometric differences between images made at different times of day and year. It also helps to moderate phenological variance and spectral errors thrown by late snowpack or early snowfall. The aim is to produce a relatively consistent set of images for comparison, while preserving strong signals of change. Because disturbance legacies usually remain detectable on a forested landscape for several years after the event (except when salvage harvest is applied), annual compositing tends to improve the accuracy of disturbance detection on net (Kennedy et al. 2010).

Finally, I computed three vegetation indices on the medoid spectral values as inputs to the temporal segmentation procedure (*§2.2.3*). Dozens of spectral indices have been proposed for distinguishing vegetation from other forms of land cover (Khorram et al. 2016). All require computations, typically on combinations of visible and infrared bands, that amplify the spectral signal of vegetative cover and diminish the signal of non-vegetative cover. Several indices have been found especially useful in identifying disturbance (Miller et al. 2009, Neigh et al. 2014b, Potter 2014, McDowell et al. 2015, Senf et al. 2015, Cohen et al. 2018). The two most often used are the Normalized Difference Vegetation Index (NDVI) (Rouse et al. 1974) and Normalized Burn Ratio (NBR) (Keeley 2009). However, recent studies have concluded that a combination of spectral indices enhances disturbance detection accuracy, likely because no one index fully captures the spectral behavior of a landscape in flux. Therefore, in keeping with recent trends toward multi-index classification, I included three indices: NDVI, NBR, and NIR_V_ (Table 2).

**Table 2.** The three vegetation indices applied in the temporal segmentation procedure, with their respective calculations and Landsat 7 Thematic Mapper (TM) band inputs.

| Index | Calculation | Landsat 7 TM Bands |
| --- | --- | --- |
| NDVI | $\frac{\text{NIR} - \text{Red}}{\text{NIR} + \text{Red}}$ | $\frac{\text{B4} - \text{B3}}{\text{B4} + \text{B3}}$ |
| NBR | $\frac{\text{NIR} - \text{SWIR}}{\text{NIR} + \text{SWIR}}$ | $\frac{\text{B4} - \text{B7}}{\text{B4} + \text{B7}}$ |
| NIR<sub>v</sub> | $\frac{\text{NIR} - \text{Red}}{\text{NIR} + \text{Red}} \times \text{NIR}$ | $\frac{\text{B4} - \text{B3}}{\text{B4} + \text{B3}} \times \text{B4}$ |

For each index, I scaled the results by 10^3^ to allow the temporal segmentation algorithm to operate on integer values without losing precision and then inverted the values so that negative index change would correspond with vegetation loss.

#### 2.2.3. Disturbance detection through temporal segmentation

The final post-processed index images were used to produce a suite of derivative change variables, which were later applied as predictors in the agent-attribution classification model.

LandTrendr (Kennedy et al. 2010, 2018) is one of several algorithms available for temporal segmentation of time series data. The core of the algorithm is an attempt to create fitted models of pixels’ spectral behavior. When configured appropriately for the image set, this process strikes a balance between removing “noisy” interannual variability while identifying the maximum possible number of significant changes in the pixel’s record. Operating sequentially on each pixel in the stack of annual medoids, the algorithm returns a series of straight-line segments joined at vertices where the change in spectral value is significant enough to be considered an inflection point. The algorithm iteratively generates simpler models and then selects the model that best fits the original data.

The Google Earth Engine implementation of the LandTrendr algorithm (Kennedy et al. 2018) was run over each of the three vegetation index collections. The codebase accepts several user-defined inputs. I constrained the analysis to starting values of NDVI > 120, NBR > 170, and NIR_V_ > 210. This trimming filtered out values that began below standard thresholds for vegetation on each index and persisted through the time series as stable non-forest. I also constrained the analysis to compute a maximum of 12 segments. I considered any change that did not persist for at least one year beyond the initial detection to be erroneous. (Fast spectral recoveries in forest remote sensing data are more often a result of radiometric noise or insufficient cloud/shadow masking than of vegetation vigor (Kennedy et al. 2010)). I therefore removed vertices where an apparent change returned to starting value within two years. For fitted model selection, I specified two best-fit criteria: the algorithm must select the model with the most vertices (again, to detect all changes in the record), but it must have a p-value within 0.75 of the model with the absolute lowest p.

Illustrations of the model fitting results appear in Fig. 4, which depicts the spectral behavior of three randomly selected pixels identified, respectively, as “Disturbed”, “Stable Forest”, and “Stable Non-Forest” through the temporal segmentation procedure. In this example, the algorithm simplified the shape of the “Disturbed” pixel’s trajectory from 17 segments in the original NIR_V_ returns to 5 in the best-fit model. The fitted model detected three disturbance events (in 2000, 2001, and 2009), followed by a period of regeneration.

**Figure 4.**
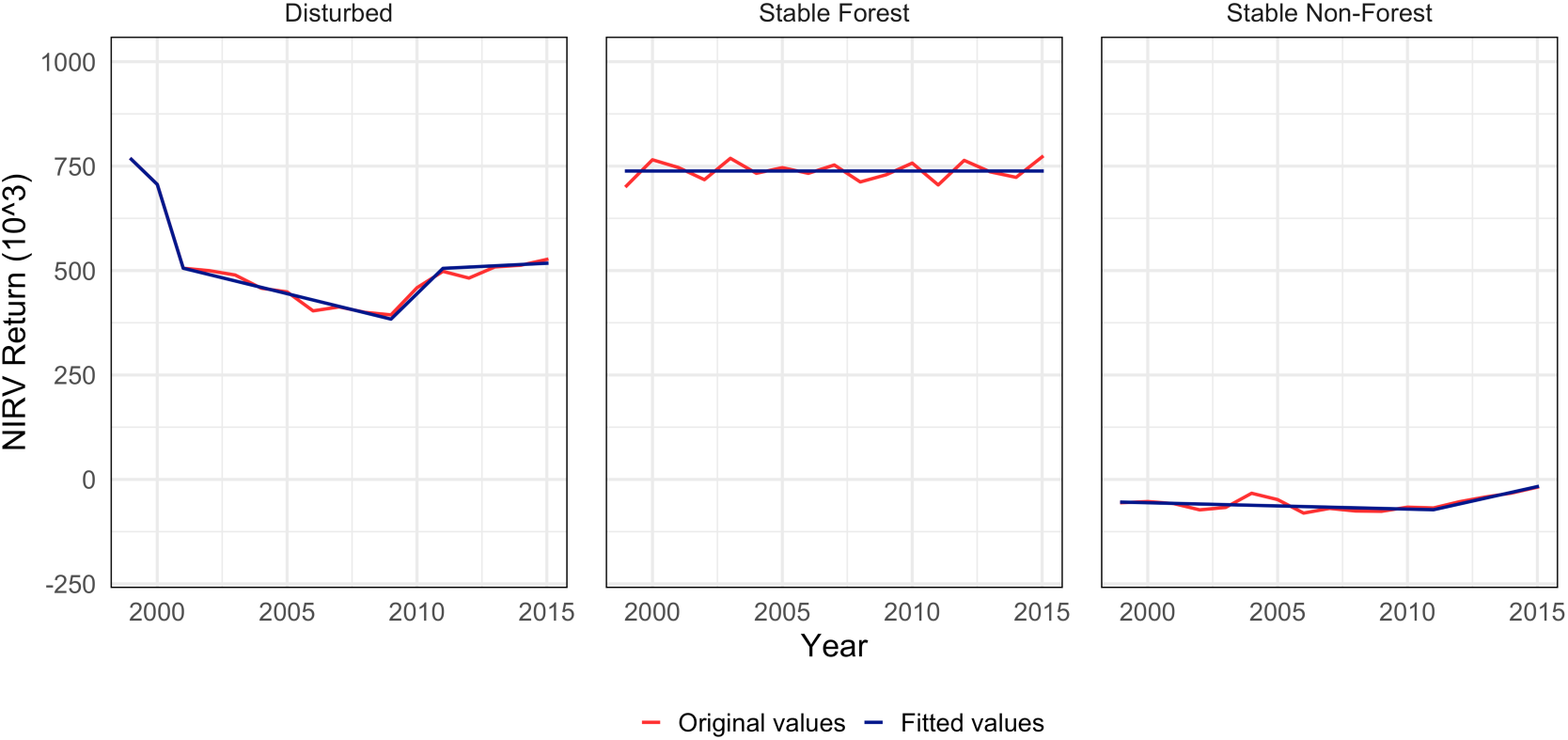
NIR_V_ and best-fit spectral trajectories of randomly selected pixels in three possible trajectory groups: “Disturbed”, “Stable Forest”, and “Stable Non-Forest.” Red lines indicate spectral trajectory based on observed NIR_V_ values. Blue lines represent the model that best simplified the trajectory shape based on thresholds defined in the temporal segmentation procedure.

The largest magnitude disturbance occurred between 2000 and 2001 (ΔNIR_V_ = 200), with a duration of one year and a rate of 200/1 = 200. The “Stable Forest” pixel’s trajectory was reduced to one segment with ΔNIR_V_ = 0. The “Stable Non-Forest” trajectory was simplified to two segments. Its absolute NIR_V_ values never exceeded the threshold for consideration as vegetation (NIR_V_ = 210), so the pixel was considered undisturbed.

After finding the best segment fits, several metrics derived from the trajectories were computed on each pixel, summarized in Table 3. The five-dimensional arrays containing these values were sliced to include only segments representing negative change greater than 10 percent, in order to remove periods of stability, periods of vegetation growth, and low-value outliers. (I make no further inferences about the excluded segments.) This process operationalized the concept of disturbance as *any negative change in the vegetation index of a pixel greater than 10 percent.* I selected the greatest-magnitude segment for each pixel. Multidimensional analysis would have exceeded the capacity of the computing resources I have available, and the greatest-magnitude disturbance on a site typically has the greatest influence on forest structure and regeneration dynamics. Finally, I cropped the arrays containing these outputs to a multipolygon shapefile delimiting the boundaries of Stanislaus National Forest (USDA Forest Service 2019).

**Table 3.**
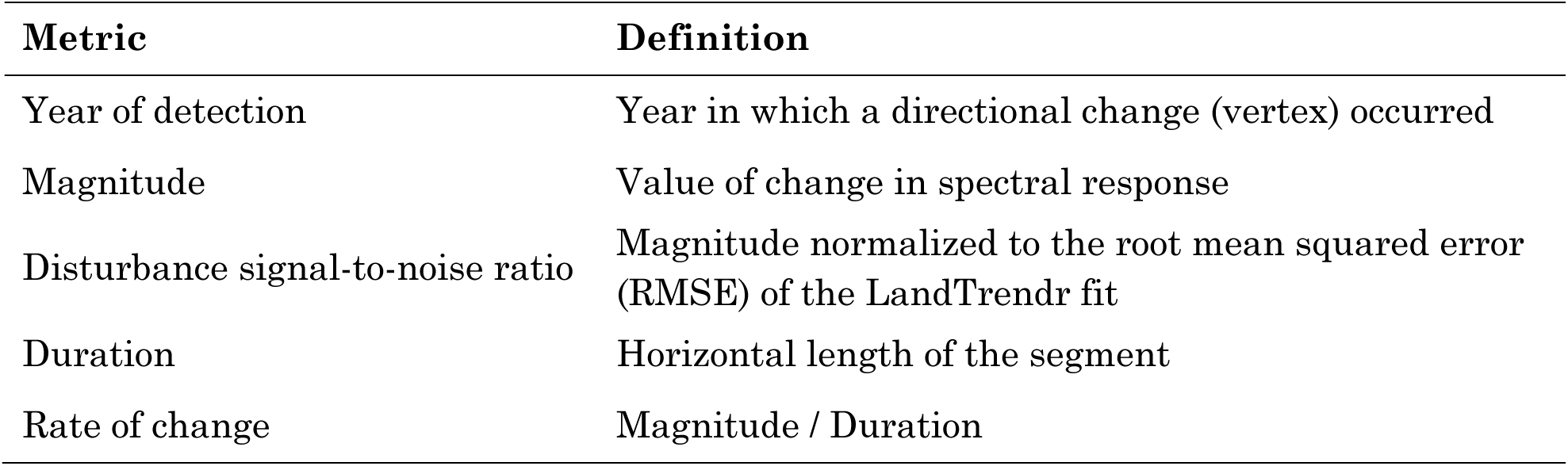
Definitions of pixel trajectory metrics: year of detection, magnitude, disturbance signal-to-noise ratio, duration, and rate. Metrics were derived through temporal segmentation of vegetation index time series.

#### 2.2.4. Derived variables

From these outputs, several derivative variables were calculated on each pixel and on pixel clusters. First, land-cover ternary maps were produced by labeling pixels according to the three possible trajectory groups identified the temporal segmentation procedure. Pixels with a detected negative change were labeled “disturbed”; undisturbed pixels with values persistently above the index vegetation thresholds were labeled “stable forest”; and undisturbed pixels with values persistently below the index vegetation thresholds as “stable non-forest.” One ternary map was created for the temporal segmentation results for each vegetation index, for a total of three maps.

Next, texture metrics were computed to quantify the textural characteristics of different disturbance classes. Using the “glcmTexture” function in Google Earth Engine (Gorelick et al. 2017), I calculated 14 GLCM metrics on each of the vegetation index images (3 indices x 17 years x 14 metrics = 294 GLCM metrics). GLCM proceeds by tallying the frequency of occurrence of pairs of pixel brightness (“grey-level”) values in a user-defined neighborhood. The frequencies are normalized to the number of observations to produce probabilities (Hall-Beyer 2017). These probabilities are then applied in a series of calculations whose results may be roughly categorized as “edge” metrics and “interior” metrics. Edge metrics produce higher values for larger and more abrupt differences in brightness values in the computing neighborhood. Interior metrics produce higher values for smaller and more heterogeneous gradients in brightness values.

Of course, edge and interior are highly scale-dependent qualities of an image, as of a landscape. In a high-resolution image of a forest, a tree crown might be discernable as an edge, while its constituent branches and leaves compose the interior; in a moderate-resolution image, only the edges of patches might be discernable, and multiple trees then make up the interior. It is important, therefore, to identify an appropriate scale for GLCM computation. Owing to the native resolution of Landsat source data and the focus on disturbance at the hectare scale or greater, I used a square 3×3 pixel window to define the computing neighborhood, so that each pixel was compared with its eight edge- and corner-adjacent neighbors in the frequency calculations. This produced texture measurements at the approximately one-hectare patch scale.

GLCM produces 14 distinct metrics, but many of them are correlated. Including them all in a classification model would produce redundancies that could reduce model skill and/or distort the evaluation of variable explanatory power (Kim et al. 2009). Based on guidance in Hall-Beyer (2017), I identified three theoretically independent measures to apply in the final analysis (Table 4).

**Table 4.**
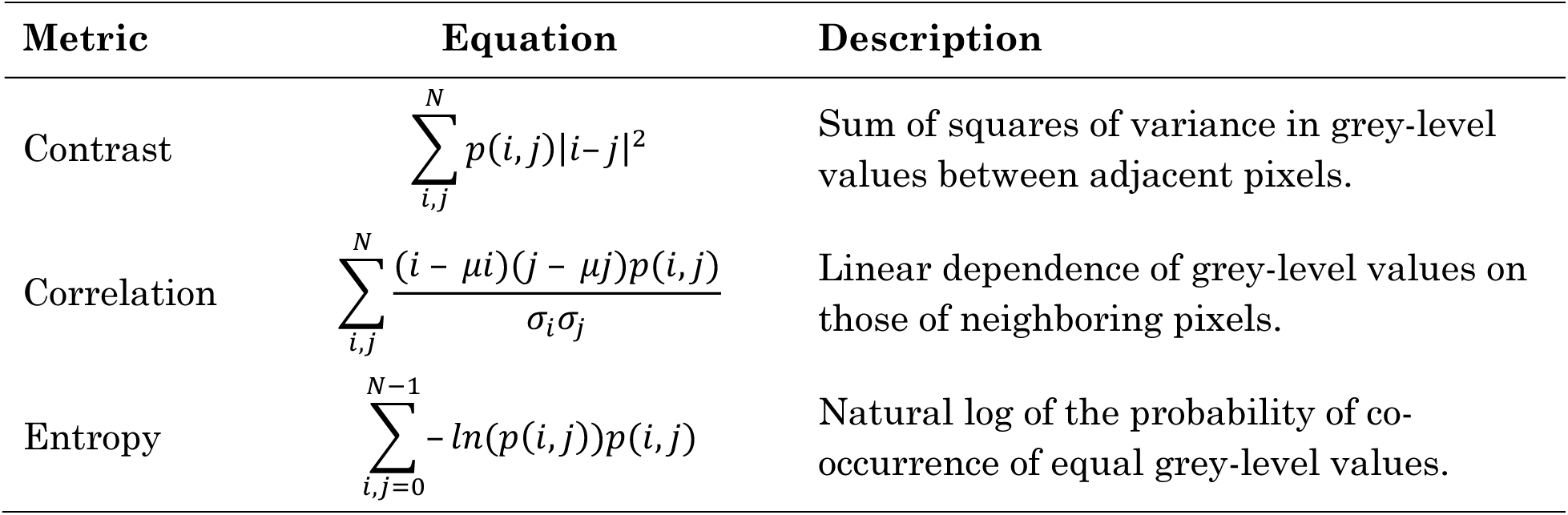
Three theoretically independent metrics for quantifying textural characteristics in forest remote sensing applications.

*Contrast* measures the intensity contrast between neighboring pixels and tends to be a reliable edge metric in vegetated landscapes (Hall-Beyer 2017). *Entropy* is also often a fruitful edge metric, particularly in areas with a high heterogeneity of radiometric intensities, as in disturbed forest with deadfall, and it may be useful for differentiating structural randomness from more uniform structures (Haralick et al. 1973). *Correlation* is an interior metric that captures the prevalence or absence of linear structure. After computing the texture metrics, I masked out undisturbed pixels and recoded them to a discrete NA value outside the NIR_V_ range.

To evaluate possible correlations between the GLCM metrics, the 14 metrics were paired separately. The Pearson correlation coefficient (*r*) was computed on all pairs and reported in a correlation matrix. The three proposed measures were confirmed for inclusion only if they were uncorrelated or weakly correlated (either *p* > 0.01 or *r* < 0.3 for significant correlations).

Next, in order to exploit the variability in geometric patterns associated with different disturbance classes, I used a 3×3 moving window segmentation algorithm to group pixels disturbed in each year into disturbed patches. I then calculated the perimeter, area, and fractal dimension of each patch. Fractal dimension is effectively an enhanced perimeter:area ratio, normalized to the expected ratio of a square and then scaled logarithmically to reduce the metric’s size dependence (*ln(0.25 * perimeter) / ln(area)*) (Turner and Gardner 2015).

#### 2.2.5. Ancillary topographical data

The final step in data preparation was to generate geophysical variables to account for topographical regulation of forest occurrence and disturbance dynamics. The National Elevation Dataset Digital Elevation Model (DEM) was resampled to 30-m pixel resolution. Slope, elevation, sine-transformed aspect and cosine-transformed aspect were computed and draped over the study site (Beers et al. 1966; Schroeder et al. 2017).

### 2.3. Random Forest classification model

A Random Forest (RF) procedure (Breiman 2001) was used to empirically model the occurrence of the four classes of disturbance identified in the reference dataset (fire, harvest, stress, conversion) and stable forest. RF is a non-parametric modeling framework that takes randomized bootstrap samples of subsets of predictor and response variables and uses them to construct an ensemble of many slightly different decision trees. When RF is used for image classification with a categorical response variable, the end-nodes of these trees comprise a set of potential classification decisions for each pixel. The procedure makes a final prediction about the correct class through a majority vote. RF was selected on three criteria. First was its ability to assimilate potentially highly correlated datastreams without overfitting and, owing to the majority-vote procedure, with only minor bias concessions. Second was its value-indifference: because the classification ultimately depends on decision trees, a variable’s relative value rather than its absolute magnitude drives the training decision. Incorporating data of widely different magnitudes does not therefore force model decisions toward predictors with higher absolute values. And third was its nearly exclusive use in other disturbance agent-attribution modeling approaches.

Two models were developed in R (R Core Team 2014) using the “ModelMap” package (Freeman et al. 2016), which optimizes ensemble modeling procedures for geospatial analysis. The first model was produced without textural variables and the second with textural variables included. In both instances, I used the 1000 points sampled from the reference dataset as trainers, with 200 points in each class. In the first model, 22 predictor variables were used; in the second model, 31 variables were used (Appendix B). The predictor sets contained true pixel values for the entire domain in the topographic and ternary variables. The remaining variables contained true values only for disturbed pixels. In these cases, the non-disturbed pixels received a discrete NA value outside their true ranges.

In early tuning of the models, different numbers of independent trees (101, 201, 501, 1001, and 2001) were tested incrementally. The number of trees required to stabilize accuracy and variable importance (i.e., the point where increases in the number of trees did not affect overall accuracy or predictor importance rank) fell between 201 and 501. The final models were set to assemble 501 trees. (The extra unit was included to break voting ties). Eight predictor variables were used per bootstrap run to decide on node splits. This was based on guidance in Freeman et al. (2016) to begin with a sample size equal to one-third the number of variables, and then to test increments above and below that number. Accuracy stabilized when eight variables were selected, so the final models were set to sample eight variables.

The predictors were randomly sampled in the construction of each tree, and all of the predictors were ultimately used. Maps of disturbance agent predictions at pixel level were produced in ModelMap.

### 2.4. Accuracy assessment and variable importance

Accuracy was assessed at three separate stages. First, the accuracy of disturbance detection in the temporal segmentation procedure (*§2.2.3*) was evaluated against a testing set (*§2.2.1*). The reference image was reduced by collapsing fire, harvest, and stress into a single “disturbed” cover class; in the resulting image, all pixels were assigned to one of three categorical values: stable forest, stable non-forest, and disturbed. A testing set was created via stratified random sampling of this image, excluding pixels that had been used in model training. The testing points were interpreted in the same manner as the training data. The procedure yielded 200 points per class, for a total of 600 testing points. These points were then used as the basis for comparison with the three cover ternary images (*§2.2.4*). Omission and commission errors were calculated for disturbed, stable forest, and unstable forest, along with overall agreement scores and Cohen’s Kappa (*κ*) statistics for each vegetation index. *κ* is a multivariate measure of accuracy that accounts for the possibility of agreement by chance. The coefficient is calculated from the error matrix and ranges from zero to one, with zero representing random-chance agreement and one representing perfect agreement. In land-cover classification, generally accepted targets for each of these metrics are overall accuracy > 85%, per-class accuracy > 70%, and *κ* > 0.61 (Foody 2002).

Second, the accuracy of disturbance detection in the RF models (*§2.3*) was evaluated against the testing set to quantify any gross information gain or loss that might have been produced in the RF. In this case, the RF maps were converted to raster format and overlaid on the ternary reference image to form a multi-band raster. The same testing points were extracted, and the same bundle of accuracy metrics was produced.

Third, the accuracy of disturbance agent-attribution was assessed using out-of-bag (OOB) estimates. OOB reports the mean prediction error for each training sample, calculated on the trees that were excluded from the bootstrap sampling operation. Because OOB observations are excluded from model training, they are thought to offer reliable accuracy estimates. Omission and commission errors, overall agreement, and *κ* statistics were measured for both RF models.

Mean decrease in accuracy (MDA) was used to evaluate the relative importance of predictors in the two models. Interpreting the absolute importance of individual predictors presents challenges in RF models because the procedure reduces hundreds of intermediate decisions to a single per-pixel vote. MDA, however, enables comparisons of relative variable importance across trees. The statistic measures how much predictive power would be lost (i.e., the percent increase in predictive error that would arise) if a variable were removed from the model. MDA values were computed and ranked for both models. Accuracy and MDA statistics were compared to evaluate the contribution of textural information to model skill.

## 3. Results

### 3.1. Texture metric correlations

Correlation testing was performed on the set of 13 GLCM texture metrics to validate the assumption that contrast, correlation, and entropy were weakly correlated and therefore contributed independent streams of information to the model. Each GLCM metric was paired separately with the other 13 in the set, and the correlation coefficient Pearson’s *r* was computed for each pair. Graphical representations of these correlations appear in Fig. 5. As expected, several of the texture metrics were closely correlated, as indicated by narrower ellipses and more-saturated colors. For instance, contrast and variance—two “edge” measures—were well correlated in the data (Pearson’s *r* = 0.975; p < 0.01). This tends to be the case when a landscape has very clearly defined edges. Where highly correlated variable pairs are thought to measure similar properties of a landscape, it is prudent to select only one member of the pair in order to develop a statistically independent metric set.

**Figure 5.**
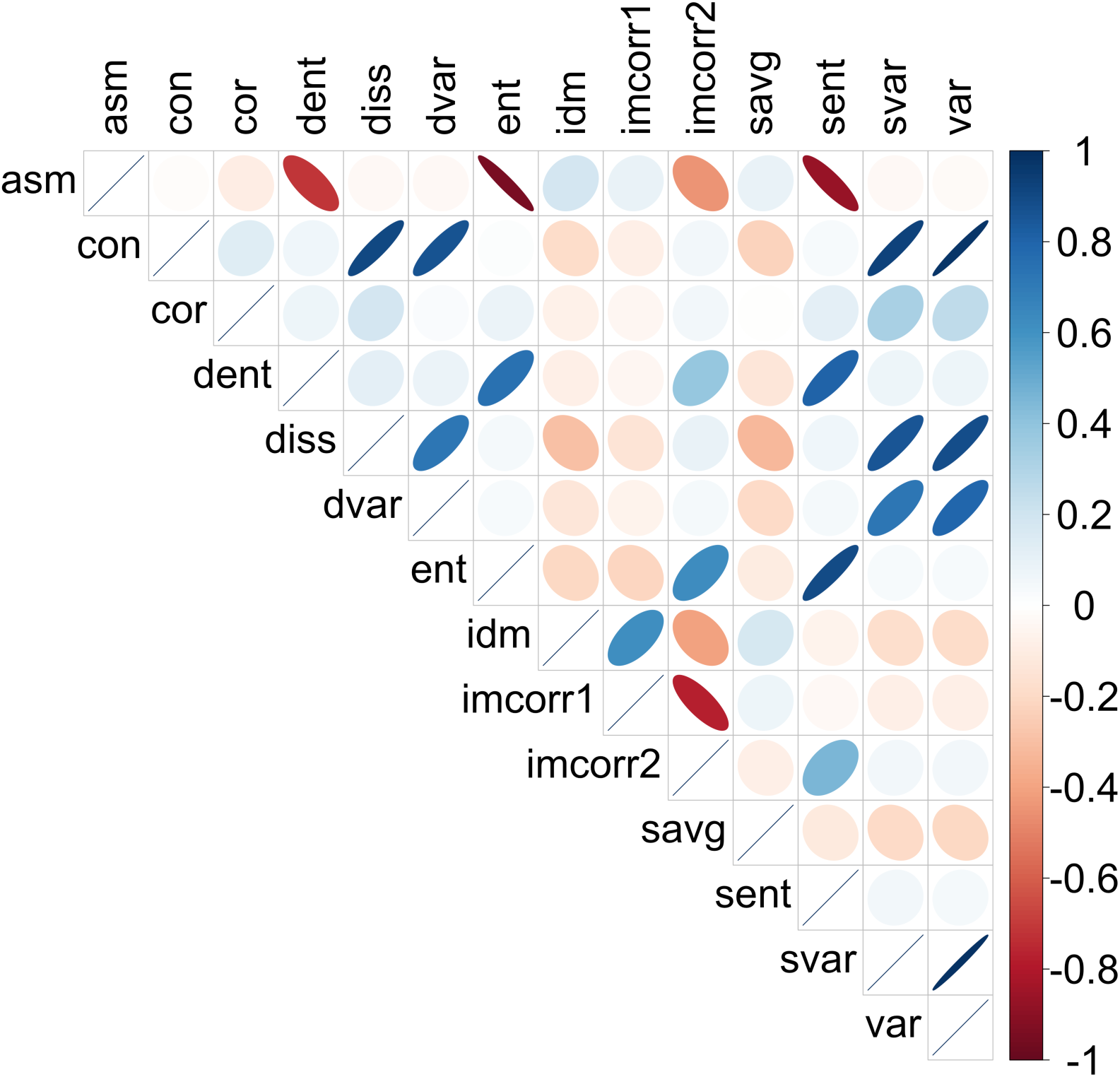
Graphical representations of the correlation coefficient Pearson’s *r* calculated for pairs of GLCM texture metrics for 1999–2015 vegetation index values. Blue values indicate positive correlation; red, negative correlation. Ellipse width and color saturation indicate the strength of the relationship. Here, the metrics contrast (con), correlation (cor), and entropy (ent) were selected for inclusion as robust independent measures of edge, interior structure, and interior randomness, respectively. Names and variable definitions for the predictor codes are in Appendix A.

The three proposed texture metrics—contrast, correlation, and entropy—were sufficiently weakly correlated to be confirmed for inclusion as textural predictors (Pearson’s *r* < 0.3) (Table 5). Spatially explicit depictions of these metrics calculated on NIR_V_ returns from 2014 are depicted in Fig. 6 for a 25km^2^ subset of the study domain.

**Figure 6.**
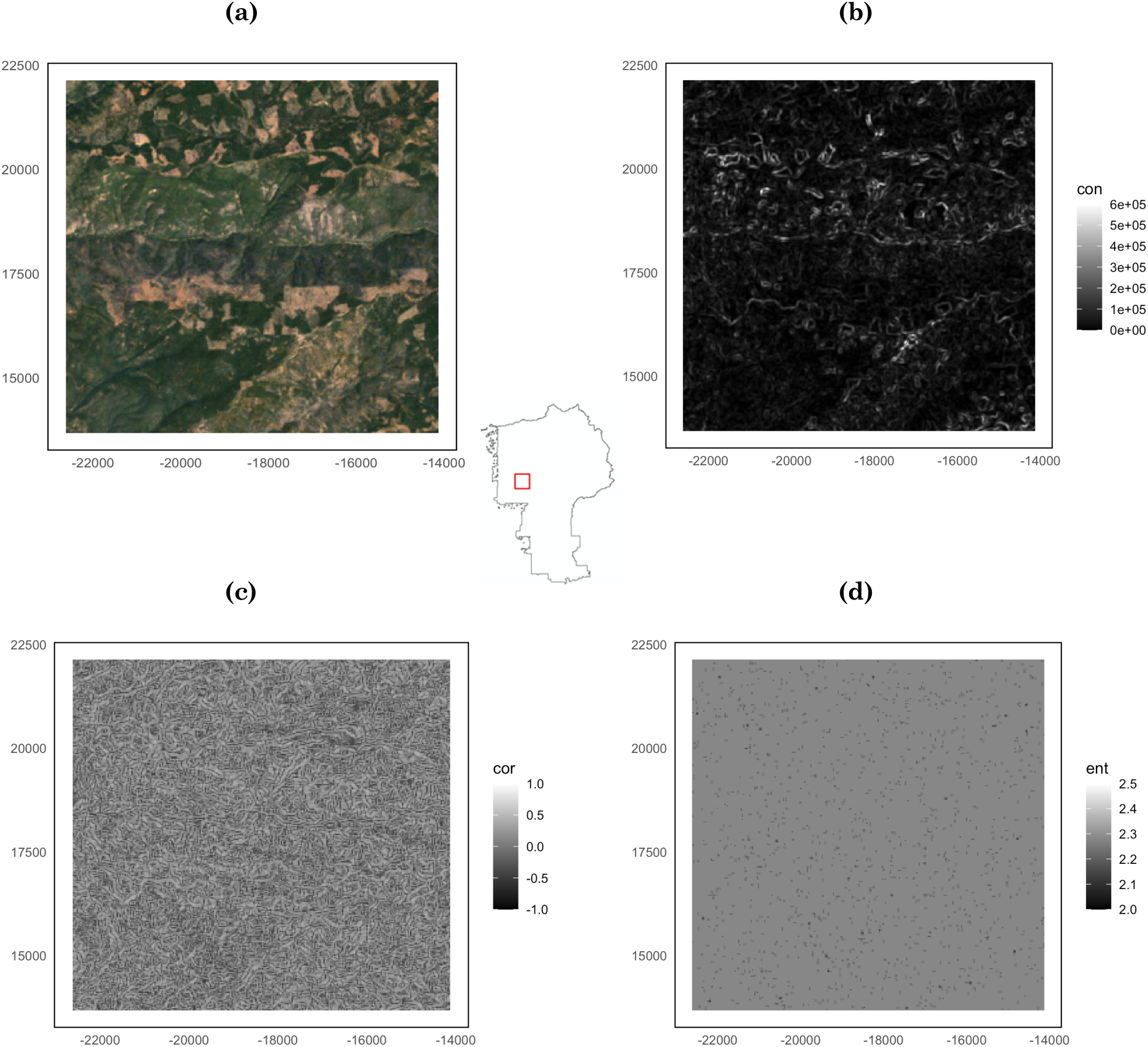
A true-color composite (a) shown alongside three GLCM texture metrics for a 25 km^2^ subset of the study domain. Contrast (b), correlation (c), and entropy (d) were calculated on NIR_V_ returns for 2014. The approximate location of the subset area within the Stanislaus National Forest boundary appears in the centered map.

**Table 5.** Pearson’s correlation coefficients for selected GLCM texture metrics.

| Variable Pair | Pearson’s *r* |
| --- | --- |
| con–cor | 0.1397*** |
| con–ent | 0.0185*** |
| cor–ent | 0.0869*** |
| *** p < 0.01 |  |

The reason for selecting independent (or weakly correlated) texture metrics has to do with how RF handles variable importance for highly correlated predictors (Schroeder et al. 2014, Freeman et al. 2015). One of the advantages of RF is that it can assimilate correlated variables without sacrificing accuracy or overfitting to the data. A tradeoff, however, is that it tends to spread out importance across those correlated variables, which makes assessing relative variable importance difficult. For the purposes of overall model accuracy, this is not such a problem. But since this study is explicitly testing the importance and contribution of distinct texture metrics, it was necessary, to the extent possible, to use independent measures.

### 3.2. Disturbance detection: temporal segmentation of vegetation index time series

The percentage of pixels identified as disturbed in the study area each year ranged widely, from 0.20% in 2011 to 15.6% in 2014 (Fig. 7). The three vegetation indices yielded similar patterns of detection, with NIR_V_ yielding the greatest total number of disturbed pixels across all years (1.45×10^6^) and NDVI the fewest (1.36×10^6^).

**Figure 7.**
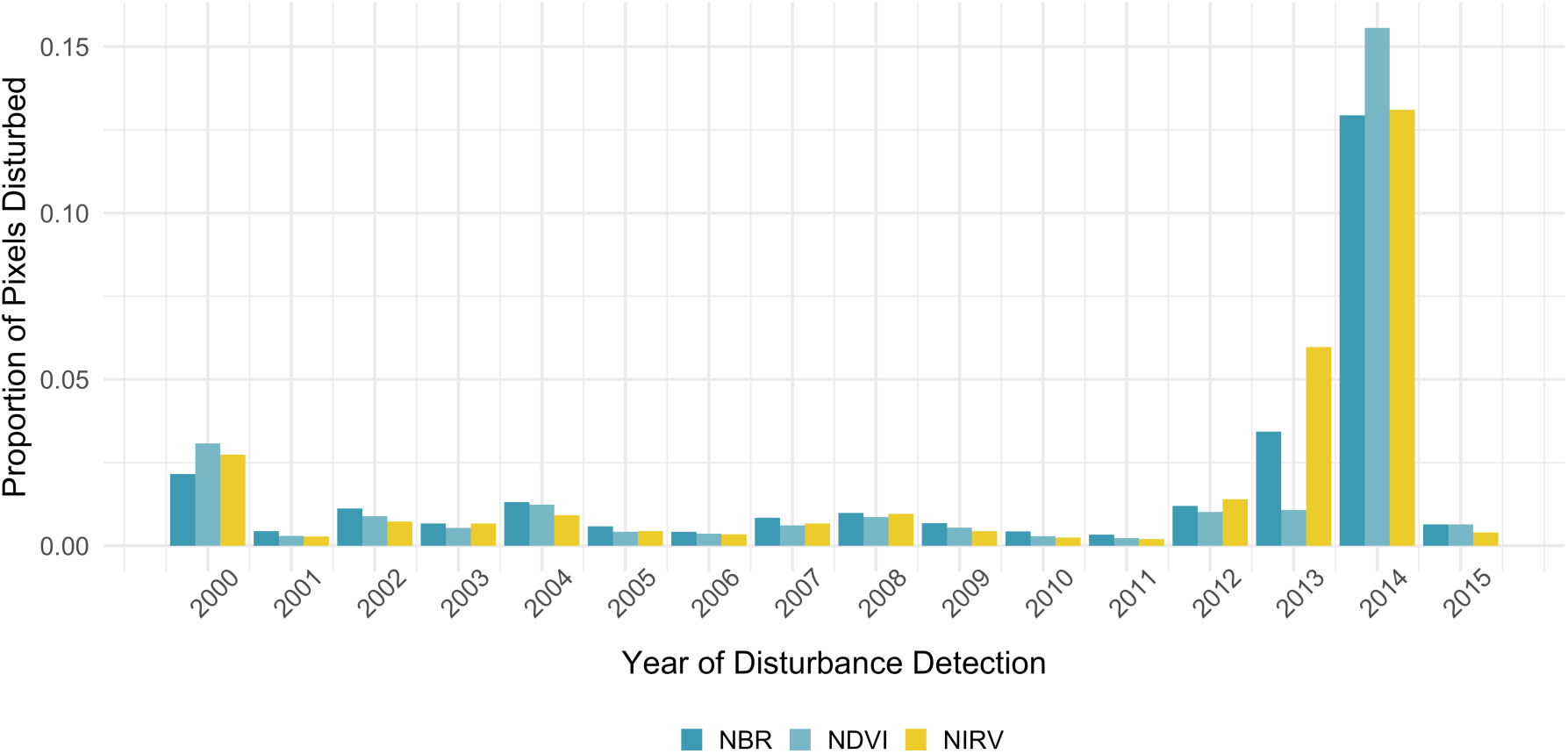
Pixels identified as disturbed as a proportion of total within Stanislaus National Forest boundaries. Disturbance was detected through a temporal segmentation procedure run on time-series stacks of NBR, NDVI, and NIR_V_ values, which were computed on annual composites of Landsat observations from 1999–2015.

The first accuracy assessment was also conducted at this stage. Overall accuracy of disturbance detection through temporal segmentation of NBR, NDVI, and NIR_V_ time-series stacks ranged from 69.3% to 74.2% (Table 6). All accuracy results were significantly better than random (p < 0.01), and there was no significant difference in accuracy among the three indices (p > 0.01).

**Table 6.** Disturbance detection accuracy and Cohen’s Kappa (*κ*) when NBR, NDVI, and NIR_V_ were assimilated separately in the temporal segmentation procedure.

| **Index** | **Accuracy** | **$\kappa$** |
| --- | --- | --- |
| NBR | 70.4% | 0.537 |
| NDVI | 74.2% | 0.598 |
| NIR<sub>v</sub> | 69.3% | 0.527 |

To enable comparisons among NDVI, NBR, and NIR_V_, their magnitude values were re-normalized to a 0 – 1 scale. RF is generally scale-indifferent, but normalizing is useful for comparing mapped values. Histograms and map renderings in Fig. 8 depict the distribution of normalized magnitudes. NDVI and NBR were similarly distributed with a mean of 0.291 and 0.266, respectively. NIR_V_ was comparatively leptokurtic, with a lower mean of 0.161.

**Figure 8.**
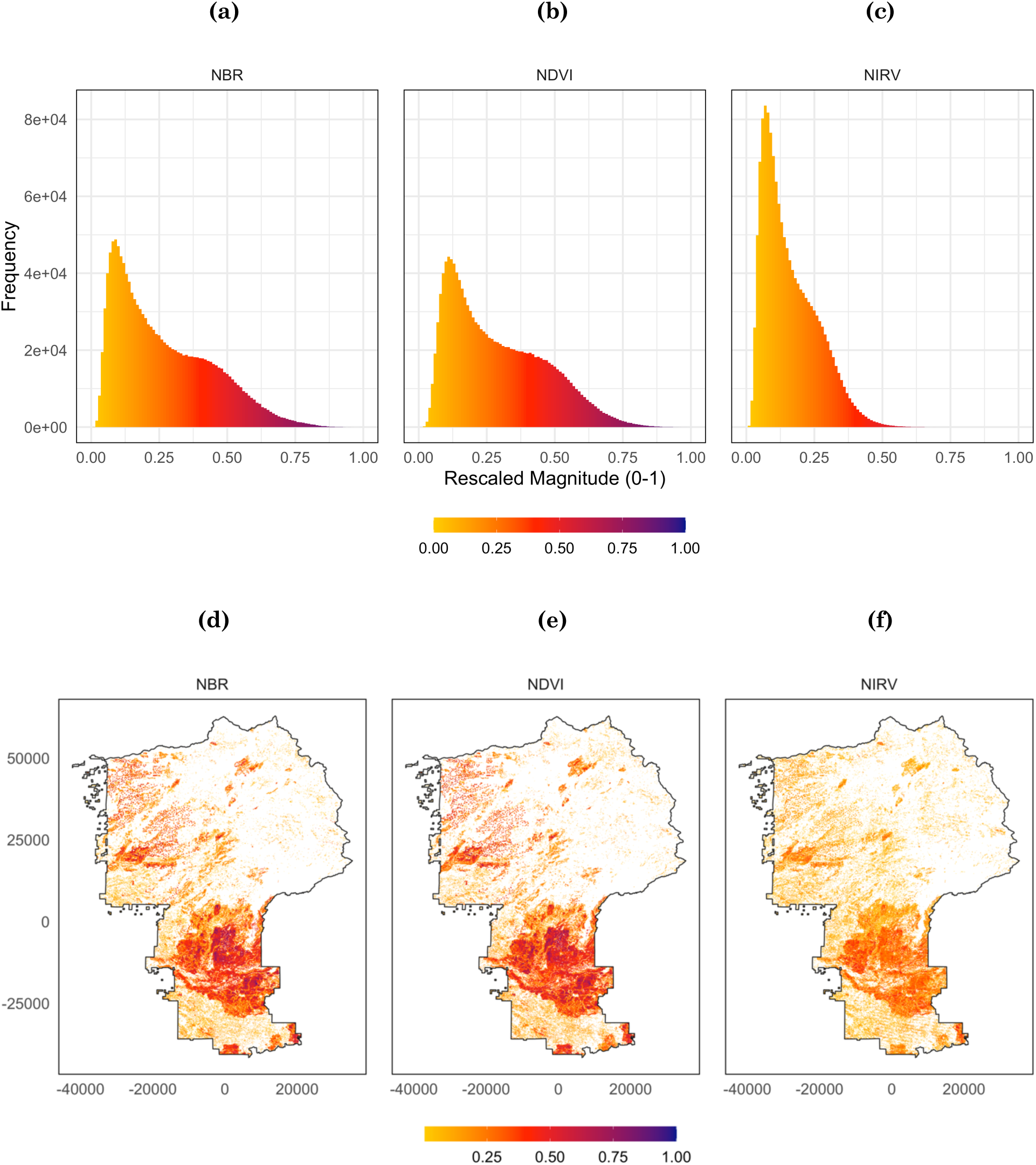
Magnitude of greatest disturbance events shown in histograms (a–c) and mapped at 30-m pixel scale (d–f) within the Stanislaus National Forest boundary. Disturbance location and magnitude were identified by temporal segmentation of NDVI, NIR_V_, and NBR time-series.

### 3.3. Distribution attribution: Random Forest classification

At this stage, the second accuracy assessment was conducted to determine whether the RF model yielded any gain or loss of skill over temporal segmentation. Accuracy was assessed in the same manner as in the first stage, except that reference points were now compared to RF-modeled images rather than the original temporal-segmentation outputs. Overall accuracy was 80.0% for Model 1 (*κ* = 0.700) and 79.8% for Model 2 (*κ* = 0.697), with no significant difference between the two (*p* > 0.01). However, RF was more adept at differentiating disturbance, stable forest, and stable non-forest than the temporal segmentation procedure alone. Overall accuracy increased with RF modeling by between 5.6% and 10.7% (p < 0.01).

Agent attribution accuracy was then assessed using the RF models’ OOB diagnostics (Table 7). The first model, which excluded textural variables, showed an overall agreement of 72.0% and *κ* = 0.650. The second model, which included the texture metrics, had an overall agreement of 72.2% and *κ* = 0.652. The difference in accuracy between the models was insignificant (p > 0.01).

**Table 7.** Error matrices for RF classification models: (a) Model 1 (texture metrics excluded) and (b) Model 2 (texture metrics included). Within the shaded box, numbers in cells represent the count of sample pixels in each category. Column values represent observations in the reference data and all sum to 200 pixels per class. Row values represent modeled agent predictions and sum to total predictions for that class. Diagonal (darker) cells contain correct identifications; off-diagonal (lighter) cells contain errors. Row and column totals, omission errors (OE), and commission errors (CE) appear in italics. Commission error is calculated as the sum of false-positive predictions (row errors) over total predictions per class. Omission error is calculated as the sum of false-negative predictions (column errors) over total reference points per class. The proportion of pixels correctly classified (PCC) appears in the bottom-right cell of each matrix.

**(a) Model 1: GLCM texture variables excluded**
|  |  | Reference |  |  |  |  |  |  |
| --- | --- | --- | --- | --- | --- | --- | --- | --- |
|  |  | Fire | Harvest | Stress | Stable forest | Stable non-forest | *Total* | *CE* |
| Predicted | Fire | 111 | 31 | 12 | 13 | 0 | *167* | *0.335* |
|  | Harvest | 36 | 105 | 4 | 56 | 0 | *201* | *0.478* |
|  | Stress | 31 | 4 | 178 | 5 | 0 | *218* | *0.183* |
|  | Stable forest | 22 | 60 | 6 | 126 | 0 | *214* | *0.411* |
|  | Stable non-forest | 0 | 0 | 0 | 0 | 200 | *200* | *0.000* |
|  | *Total* | *200* | *200* | *200* | *200* | *200* | *1000* | *PCC* |
|  | *OE* | *0.445* | *0.475* | *0.110* | *0.370* | *0.000* | *PCC* | ***0.720*** |

(b) Model 2: texture variables included
|  |  | Reference |  |  |  |  |  |  |
| --- | --- | --- | --- | --- | --- | --- | --- | --- |
|  |  | Fire | Harvest | Stress | Stable forest | Stable non-forest | Total | CE |
| Predicted | Fire | 110 | 28 | 10 | 14 | 0 | 162 | 0.321 |
|  | Harvest | 36 | 120 | 7 | 70 | 0 | 233 | 0.485 |
|  | Stress | 35 | 5 | 179 | 3 | 0 | 222 | 0.194 |
|  | Stable forest | 19 | 47 | 4 | 113 | 0 | 183 | 0.383 |
|  | Stable non-forest | 0 | 0 | 0 | 0 | 200 | 200 | 0.000 |
|  | Total | 200 | 200 | 200 | 200 | 200 | 1000 | PCC |
|  | OE | 0.450 | 0.400 | 0.105 | 0.435 | 0.000 | PCC | 0.722 |

Omission and commission errors from both models indicated that stress and stable non-forest had the highest model agreement with reference data (refer to the “CE” column and “OE” row in Table 7). Model errors for these classes were relatively well balanced—OE and CE scores fell within ±10% of each other—which suggests that the model was appropriately tuned to the data. Stable forest and harvest were also well balanced, but their errors were higher (> 35.0%), and they were systematically confused with one another (60 instances in Table 7a and 47 instances in Table 7b). Fire’s accuracy was moderate (balanced accuracy = 79.8% in Model 1 and 76.0% in Model 2), but it was frequently confused with all other classes except non-forest, as the false-positive and false-negative entries along the “Fire” row and column indicate.

### 3.4. Predictor variable importance

Table 8 reports predictor variable importance in terms of the decrease in overall model accuracy (mean decrease in accuracy, MDA) that would result if a given variable were excluded from the model. The three most powerful predictors in both models were the land-cover ternary generated from NDVI segmentation, elevation, and fractal dimension computed on NDVI. Fractal dimension from all three vegetation indices emerged in the top 15 explainers in both models. In the second model, the texture metrics appeared to promote the relative importance of slope. While texture metrics added comparatively little explanatory power, entropy was the highest-ranked contributor of the texture metrics.

**Table 8.** Relative importance of the top 15 predictor variables in Model 1 (textures excluded) and Model 2 (textures included). Importance is expressed in terms of mean decrease in accuracy (MDA), the accuracy penalty that would result if a variable were excluded from the set of predictors. Texture variables that appeared in the top 15 for Model 2 are in **bold** type.

| Model 1: texture metrics excluded |  |  | Model 2: texture metrics included |  |
| --- | --- | --- | --- | --- |
| Rank | Variable | MDA | Variable | MDA |
| 1 | NDVI ternary | 67.9 | NDVI ternary | 39.2 |
| 2 | Elevation | 53.2 | Elevation | 37.1 |
| 3 | NDVI fractal dimension | 35.1 | NDVI fractal dimension | 24.0 |
| 4 | NDVI disturbance rate | 28.7 | Slope | 21.7 |
| 5 | Slope | 25.4 | NIR<sub>v</sub> ternary | 20.8 |
| 6 | NIR<sub>v</sub> fractal dimension | 21.2 | NDVI disturbance magnitude | 17.9 |
| 7 | NDVI disturbance magnitude | 20.5 | NDVI disturbance rate | 17.7 |
| 8 | NIR<sub>v</sub> ternary | 19.2 | NDVI disturbance year | 17.6 |
| 9 | NBR disturbance rate | 18.9 | NIR<sub>v</sub> fractal dimension | 15.6 |
| 10 | NDVI disturbance year | 18.4 | **NDVI entropy** | **15.4** |
| 11 | NIR<sub>v</sub> disturbance magnitude | 16.9 | NBR ternary | 15.0 |
| 12 | NBR fractal dimension | 16.5 | NBR fractal dimension | 14.3 |
| 13 | NBR disturbance magnitude | 15.2 | **NIR<sub>v</sub> contrast** | **14.2** |
| 14 | NBR dsnr | 14.6 | NIR<sub>v</sub> disturbance rate | 13.9 |
| 15 | NIR<sub>v</sub> disturbance rate | 14.2 | **NIR<sub>v</sub> entropy** | **13.9** |

Notably, in Model 2, the absolute values of predictor MDA decrease for all variables, despite similar rankings. This suggests that textures are not simply “noisy” predictors but contribute information to the classification decision; they also appear to balance the overall distribution of importance across predictors.

#### Spatial predictions of disturbance agents

The predicted forest disturbance agents were mapped alongside stable forest and non-forest predictions (Fig. 9). The maps confirm that the models were able to distinguish effectively among disturbance agent classes.

**Figure 9.**
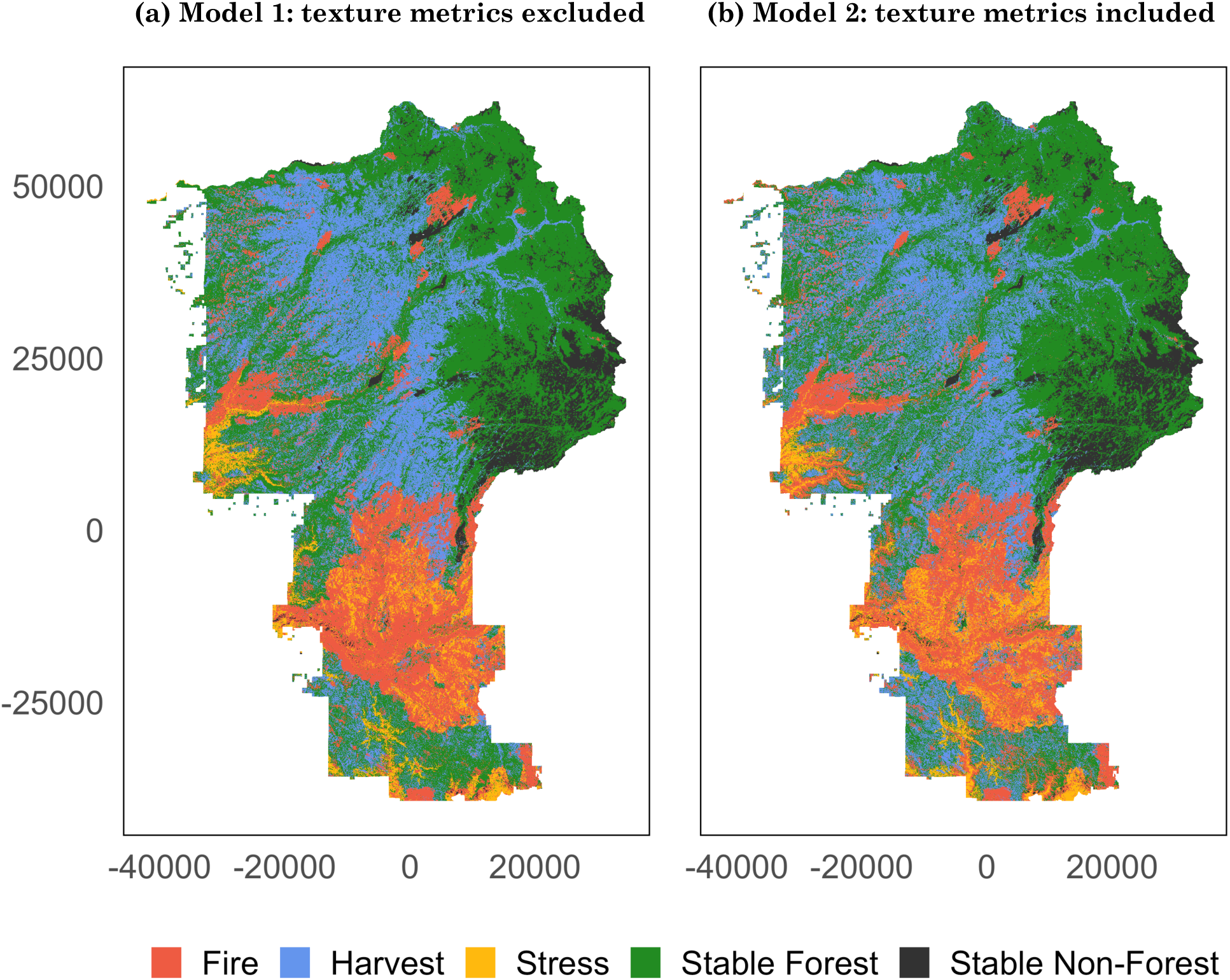
Mapped predictions of disturbance agents: (a) Model 1 (texture metrics excluded) and (b) Model 2 (texture metrics included).

The models were particularly sensitive to differences between fire and stress, which tended to co-occur at lower elevations. However, the error matrices revealed high rates of false-positive stress identification; this occurred mostly around the margins of fire perimeters, suggesting that the models may confuse stress with low-intensity fire. In general, mapped predictions of fire were well resolved and agreed closely with CalFire FRAP perimeters.

The overprediction of harvest identified in the error matrices bears out in the maps. In reality, harvest is generally constrained between 1000–1500m. Harvest does appear less frequently in the northern and eastern sections of the maps; these unharvested areas closely match the Mokelumne, Carson-Iceberg, and Emigrant Wilderness boundaries, where harvest is proscribed. These areas also occur at higher elevations, which points to the strong effect of elevation in the models. Notable exceptions are the distinctive narrow stretches identified as harvest in the easternmost portions of the maps. These follow the Clark and Middle Forks of the Stanislaus River and are contained within Carson-Iceberg Wilderness. There is no record of harvest in these areas in the LANDFIRE reference data.

Among the predicted disturbance agent classes, harvest was the most prevalent, followed by fire and stress (Fig. 10). Forest persisted in more than 40% of the area over the study period. Because the analysis is temporally indifferent and does not account for regeneration, any pixel identified as disturbed retains this status, regardless of when the disturbance was detected.

**Figure 10.**
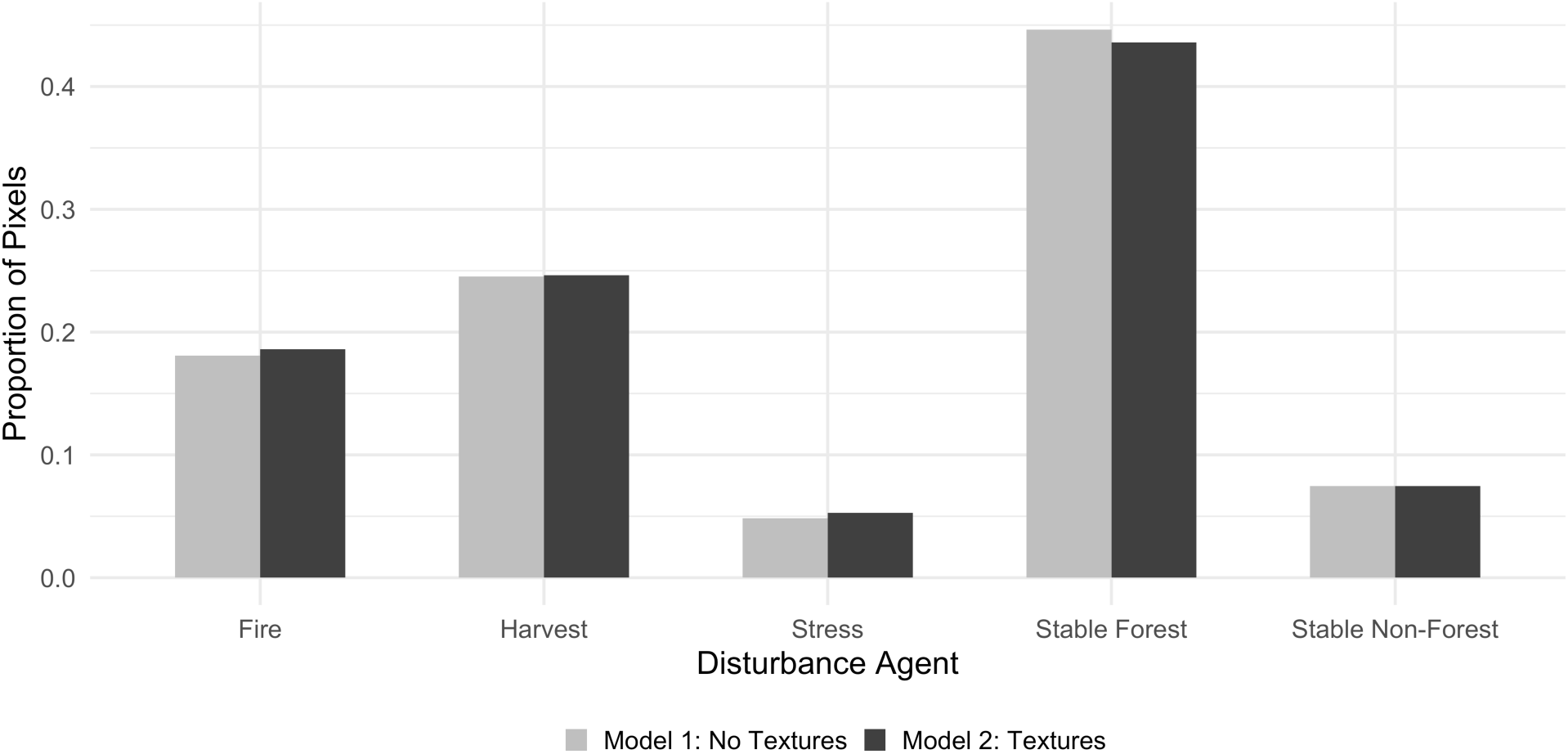
Proportion of pixels in modeled results by predicted disturbance agent or stable status.

Because elevation proved to be a strong predictor, I further analyzed the relationship between elevation and disturbance agent prevalence (Fig. 11). Stable forest was widely distributed across elevations up to ∼2800m, and non-forest was self-evidently concentrated at altitudes above 2000m (i.e., exposed granite batholith and alpine vegetation communities). Harvest appeared to be more constrained to mid-elevations, while fire and stress tended to co-occur at lower elevations, as is evident in the varying means, *x*-widths and *y*-densities of the violin plots.

**Figure 11.**
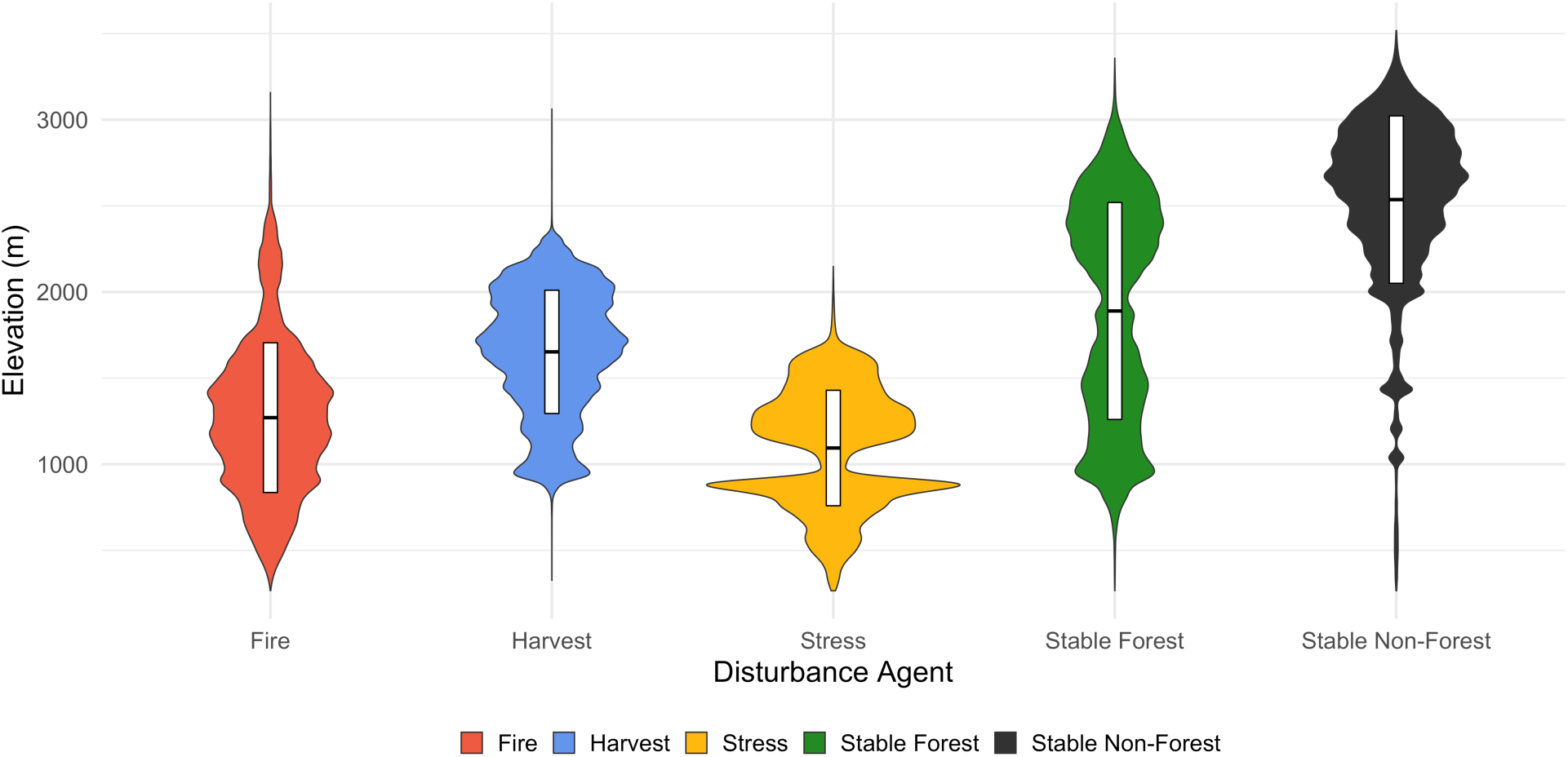
Violin plots depict elevational regulation of different disturbance agents and forest cover. “Wider” oblongs indicate more peaked distributions in one or more elevational bands, while “taller” oblongs indicate a more uniform distribution along the elevational gradient. Fire and stress appear to co-occur at lower elevations, while harvest is concentrated in mid-elevations. The white boxes in the centers of the oblongs depict the median and interquartile range of elevation.

## 4. Discussion

### 4.1. Conceptual challenges

As patterns of forest disturbance continue to shift in the Sierra Nevada of California, it is imperative that forest monitoring programs efficiently and accurately resolve not only the spatial and temporal qualities of disturbance events, but also their causes. Disturbance agents have variable impacts on forest composition, structure, and function, and effective forest management will increasingly depend on robust estimates of prior disturbance-agent prevalence as well as skillful predictions of future trends. An ideal product to satisfy this need would be an accurate, full-coverage map of historical disturbance that (a) renders the events explicitly in space and time, (b) accounts for their drivers, and (c) can be readily produced and updated with minimal analyst oversight. Achieving such an ideal through a modeling approach requires overcoming at least three major challenges. First is the basic difficulty of differentiating change agents (Kennedy et al. 2015). Disturbance is not inherently related to the spectral signals captured by most remote sensors, and different agents of disturbance can leave identical spectral signatures. While spectral information contributes substantially to disturbance detection (Cohen et al. 2018) and goes part of the way toward agent attribution (Schroeder et al. 2017, Shimizu et al. 2019b), additional information about the landscape and the processes manifest on it is necessary for a reliable and generalizable approach. A second challenge is spatial scale. Remote sensing operates at the scale of the pixel and is for the most part limited by the native resolution of satellite and aerial sensors (although the increasing availability of very high-resolution images and the promising development of new data fusion methods are rapidly diminishing the size of this challenge) (Cakir et al. 2006, Khorram et al. 2016). Ecological processes occur at scales much smaller and much larger than the 30-m pixel used in this study. Disturbances affect individual trees, and they affect entire landscapes. Reliance on pixel scale means accepting error at both ends: in generalization of sub-pixel information and in over-specification of behavior that is in fact occurring across aggregations of pixels. The third challenge has to do with heterogeneity in the spatial extent, temporal duration, and intensity of disturbances’ impacts on vegetation. Variability in harvest densities, for instance, yields considerable heterogeneity within what would ideally be considered a uniform category of change agent. The same can be said for variable-density thinning treatments, species-selective beetle kill, and fire. Here, I have described an approach that incrementally advances the field toward addressing these challenges.

### 4.2. Disturbance detection

In simply detecting disturbances, the RF model performed better than the temporal segmentation procedures run on NDVI, NBR, and NIR_V_ time-series stacks. This additional improvement from RF was likely the result of combining information from multiple spectral indices. Indeed, this was consistent with recent findings in the literature that combining multiple indices can yield higher detection accuracy (Kennedy et al. 2015, Schroeder et al. 2017). The hypothesized reason for this effect is that no single index accounts for the full range of spectral behavior in a disturbed forested landscape. The fact that detection accuracy was not significantly different when any single index was used further confirms this inference.

Given these results, NIR_V_ did not appear to aid disturbance detection on its own. It did not appear to detract either, although the indices were not tested systematically against one another. NIR_V_ captured a broader range of vegetation changes than NDVI and NBR, based on its greater total identification of disturbed pixels. Higher peaks and positive skew in the distributions of raw NIR_V_ values and ΔNIR_V_ values suggested that NIR_V_ was more sensitive to subtler negative changes in vegetation than the other two indices were. However, this behavior may have also been driven by the NIR multiplier in the NIR_V_ calculation or by noisy false-positive detection.

In any case, the total number of disturbed pixels identified across the three indices (Fig. 7) appeared to vary more consistently with year of detection than with index. While NBR’s disturbed total consistently exceeded that of the other two indices in low-disturbance years, NDVI was anomalously high in 2014 and anomalously low in 2013. NIR_V_ total disturbed was anomalously high in 2013. No ready pattern emerges from this behavior. However, 2013 and 2014 witnessed the Rim Fire, represented in Fig. 9 by the large swath of fire-attributed pixels in the southern third of the maps. This was immediately predated by intense drought-related desiccation stress in 2012–2013. It may be the case that NIR_V_ is more sensitive to stress responses, while NDVI is more sensitive to fire responses. On this interpretation, NBR’s moderate detection of fire may be more accurate. The Rim Fire years notwithstanding, there was no obvious increasing or decreasing trend in total disturbance evident over time, though a discernible trend would not necessarily be expected on a 16- year timescale.

### 4.3. Agent attribution

The procedure appeared to capture the broad categories of disturbance operating in Stanislaus National Forest between 1999 and 2015. The results underscore the need to incorporate data beyond first-order spectral-reflectance metrics. Two measures of landscape position, elevation and slope, ranked among the top five predictors in both models. Their relative importance is most likely a consequence of how these topographical characteristics regulate the presence and structure of vegetation. Topography also influences disturbance processes: harvest tends to occur at lower elevations and on shallower slopes; fire has been found to spread more rapidly on steeply inclining slopes and to burn more intensely on steeply declining slopes. Some of the beetle infestations of the early 2010s also occurred within distinct elevation bands, partially a result of elevational controls on tree species distributions. Fractal dimension is a landscape shape metric several processing steps removed from raw spectral returns, yet it was the third strongest explainer in both models. This hints at the importance of scale in this modeling approach; fractal dimension exploits the sizes and shapes of disturbed patches, while other predictors in the set primarily act at the pixel scale. Spatial extent is a key characteristic of disturbance legacies and is frequently differentiable by agent on the ground. Its appearance as one of the more important predictors squares with this observation.

The overall skill of the model, evaluated in terms of model accuracy (∼72%) was reasonable, but not exceptional. Per-class accuracy ranges between 71% and 100% were on par with the those in the most successful agent-attribution models in the literature (Kennedy et al. 2015, Schroeder et al. 2017, Shimizu et al. 2019a). Those studies yielded higher overall accuracy values than the method in this paper (78–95%). Their *κ* statistics ranged between 0.40 and 0.85. In the Schroeder et al. (2017) study, the scene that overlapped Stanislaus National Forest actually returned the highest accuracy rate (95%) of all of the scenes in their investigation. My results were significantly less robust, despite similar agent-class groupings, reference data, and input variables in the texture-free model. One reason for the discrepancy could be that Schroeder et al.’s time series ended in 2010, before the major drought and Rim Fire; their observations may have included less stress-related spectral change overall, which may have dampened confusion of stable forest, harvest, and stress. Another major divergence was that they used VCT for temporal segmentation; it would be worthwhile to test the impacts of assimilating VCT-derived vs Landtrendr-derived disturbance metrics for agent attribution in the future.

At the class level, considerable confusion arose between stable forest and harvest, resulting in systematic overprediction of harvest. Commission error for harvest exceeded 0.45 in both models. The confusion here likely results from different mechanical harvest treatments being compressed into one category. Selective removal and thinning were grouped together with clear-cuts, a decision that likely expanded the dimensional space for harvest enough that it caused model votes for harvest to also capture stable forest. The balanced omission and commission errors for these two classes is a good indicator that this was the case. A second source of error may have been the masking of stable forest pixels in several of the predictors (i.e., magnitude, year of detection, rate, fractal dimension, and the three texture metrics). Masking was the best solution to an intractable dilemma: using full coverage data for those metrics would have entailed assimilating a separate image for each year. For the texture metrics alone, this would have yielded 153 distinct images (3 variables x 3 indices x 17 years), a rate of expansion that would have quickly exhausted available computing capacity and likely would have biased the model toward the orders-of-magnitude more prevalent variable types. In fact, in early iterations of the model, I tested this possibility using full-coverage annual textures for NIR_V_ alone. The model skill was insignificantly different from the model described in this paper. And although textures did contribute a greater share of predictive power, this likely had more to do with their dominance of the share of predictors.

### 4.4. Spatial patterns and prospects for application

With the exception of overpredicted harvest, the location and distribution of attributed change agents cohered with expectations for the study site, from the minimum-mapping unit scale of one hectare up to the full National Forest scale. The fact that reasonably accurate disturbance-agent predictions can be made with a very small proportion of pixels used as training points (0.07% of total) underscores the promiseof thise approach for reducing the time and resource requirements of agent-explicit disturbance detection at the landscape scale.

### 4.5. Contribution of texture metrics

Textures contributed not at all to the absolute accuracy of the models and only negligibly in terms of the relative importance of predictors. The insignificant results mean that the null hypothesis cannot be rejected, and that textures have little effect on the modeling method’s predictive skill. Several inferences seem plausible. The first is that textural information may straightforwardly fail to add power to differentiate among disturbance legacies. This would seem to be confirmed by the null difference in overall model accuracy. A second interpretation is that textural information contributes to skill, but it is a much weaker explainer than the topographic and shape variables that drive most of the prediction. This would seem to be confirmed by the appearance of the NDVI entropy metric among the top ten predictors in the second model.

One important limitation confronts interpretation of individual predictor importance. Because of RF’s tendency to distribute importance across correlated variables, retaining correlates in the set will also influence the relative importance of independent metrics. Several of the variables were correlated; most of those derived directly from temporal segmentation (i.e., the “Disturbance” category in Appendix B) had paired Pearson’s *r* coefficients > 0.50 (p < 0.01). On the whole, the texture metrics were not well correlated with any other variables (*r* < 0.30, p < 0.01). One exception was entropy, which varied with all of the “Disturbance” variables (*r* > 0.50, p < 0.01).

In the course of this study, I was unable to adjust satisfactorily for this distortion. In prior disturbance agent attribution studies, authors have either ignored the variable interdependence problem or computed a rank sum of importance for groupings of correlated variables (Schroeder et al. 2014); this requires observations from multiple independent model replicates and so was infeasible for this single-domain study. Another solution might be to systematically remove variables from correlated pairs. However, exploratory tests of this approach significantly reduced model skill and so were rejected for this project. A third option could be to use factor analysis to compress the variable set into a smaller collection of uncorrelated factors and to rank this smaller collection according to a sum or mean rule. This seems like a promising direction, but acquiring an honest operational understanding of factor analysis exceeded the scope of an already capacious project.

In sum, while care was taken to identify independent measures of texture in order to evaluate their importance in comparison with one another, inferences about any variable’s overall rank in the predictor set may be distorted by interdependences among other variables. Accordingly, there are limits to the inferences that can be drawn from variable importance.

It may be the case that landscape textures are important for discriminating disturbance legacies, but that texture was insufficiently operationalized in this study. One potential weakness was the aforementioned masking of stable forest and stable non-forest in several of the predictors. In future work, it would be advisable to study a smaller area over a shorter timescale, focusing on pixels where only stable forest and harvest co-occur. Including full-coverage spectral and textural metrics in this case could improve model skill markedly.

Another underexplored area is the spatial scale of texture computation. The ability of edge and interior texture metrics to differentiate disturbance agents is necessarily a function of the scale at which disturbance occurs. Calculated in a 3×3 pixel neighborhood, contrast was robust to edges of harvested and stressed patches (Fig. 6). Correlation and entropy were less adept at discriminating among interior behaviors in disturbed and stable patches. A promising direction for further study would be to evaluate a wider range of neighborhood sizes. Including 24-neighbor and 224-neighbor iterations, for example, might help to identify interior patch structures that aren’t detectable in an eight-neighbor window. This information could enhance the contribution of textural metrics.

Finally, a major unresolved issue for this study and other agent-attribution approaches is the lack of an external reference dataset with sufficient temporal and spatial resolution across the length of the Landsat record to use for independent model training and validation. This is something of a chicken-and-egg problem. Using incomplete ancillary datasets and records to manually verify disturbance occurrence and agent class for 1000 training points is a tedious and error-prone exercise that further underscores the need for a more reliable modeling approach. But in the absence of a valid independent reference, a generalizable modeling approach remains difficult to achieve. A randomly sampled and verified set of retrospective disturbance points with error terms would help to an extent. However, because of the conceptual fuzziness of ecological disturbance noted in the introduction to this paper, an absolute reference may be inherently elusive, especially for agent classes that are difficult to differentiate even through on-the-ground study, such as drought stress and beetle kill. In light of this constraint, most agent-attribution approaches have aimed not for perfect classification agreement, but for improvement over the inconsistent and discontinuous data products currently in widespread application in forest management. Acknowledging that the approach described here inherits some uncertainty from reference data, remotely sensed data, and model decisions alike, it still succeeds on this more modest criterion of incremental improvement.

## 5. Conclusions

The objective of this project was to develop and test an integrated empirical modeling method for attributing forest disturbances to particular agents. The motivation was twofold: to advance a burgeoning field of methodological inquiry in the remote sensing of forest resources and to enhance the information streams available to resource and conservation managers for decision-making regarding disturbance adaptation and mitigation. The approach presented here satisfies both. The method yields adequate identification of disturbance location and moderate attribution accuracy for multiple disturbance agents. While texture as it was operationalized here did not meaningfully contribute to model skill, the results further confirm that information beyond spectral reflectance records is required for accurate agent attribution. As a proof-of-concept, this study offers a strong foundation for future work, which should focus on improving the overall efficacy of the models and generalizing them for systems beyond the Central Sierra Nevada.

## Appendix A. GLCM texture variable definitions

**Table A.**
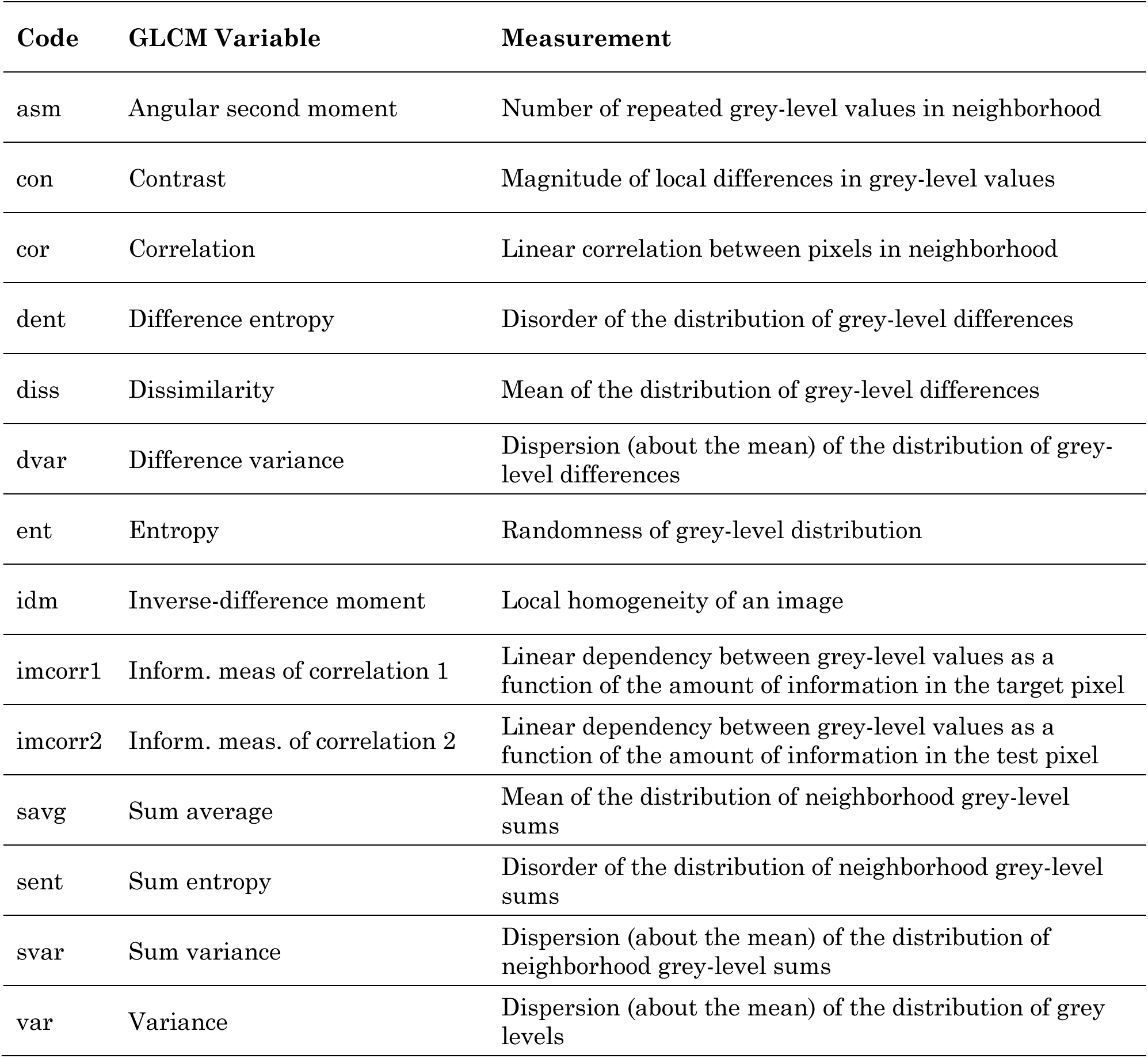
Names and definitions of the 14 GLCM texture metrics calculated on annual NDVI, NBR, and NIR_V_ composites from 1999–2015. These metrics were tested for correlation using Pearson’s *r*, and the statistics were reported in a correlation matrix (Fig. 5 in the main text). Definitions are from Zwanenburg et al. (2016), Gorelick et al. (2017), and Hall-Beyer (2017).

## Appendix B. Model predictors

**Table B.1.** Predictors for Model 1: textures excluded

| Category | Variable | Name | Coverage |
| --- | --- | --- | --- |
| Topographic | Elevation | elevation | Full domain |
|  | Cosine-transformed aspect | cos_aspect |  |
|  | Sine-transformed aspect | sin_aspect |  |
|  | Slope | slope |  |
| Ternary | NBR cover ternary | NBR_coverTernary | Full domain |
|  | NDVI cover ternary | NDVI_coverTernary |  |
|  | NIRv cover ternary | NIRV_coverTernary |  |
| Shape | NBR fractal dimension | NBR_FracDim | Disturbed pixels |
|  | NDVI fractal dimension | NDVI_FracDim |  |
|  | NIRv fractal dimension | NIRV_FracDim |  |
| Disturbance | NBR disturbance magnitude | NBR_mag | Disturbed pixels |
|  | NBR disturbance year of detection | NBR_yod |  |
|  | NBR disturbance signal-to-noise ratio | NBR_dsnr |  |
|  | NBR disturbance rate | NBR_rate |  |
|  | NDVI disturbance magnitude | NDVI_mag |  |
|  | NDVI disturbance year of detection | NDVI_yod |  |
|  | NDVI disturbance signal-to-noise ratio | NDVI_dsnr |  |
|  | NDVI disturbance rate | NDVI_rate |  |
|  | NIRv disturbance magnitude | NIRV_mag |  |
|  | NIRv disturbance year of detection | NIRV_yod |  |
|  | NIRv disturbance signal-to-noise ratio | NIRV_dsnr |  |
|  | NIRv disturbance rate | NIRV_rate |  |

**Table B.2.** Predictors for Model 2: textures included

| Category | Variable | Name | Coverage |
| --- | --- | --- | --- |
| Topographic | Elevation | elevation | Full domain |
|  | Cosine-transformed aspect | cos_aspect |  |
|  | Sine-transformed aspect | sin_aspect |  |
|  | Slope | slope |  |
| Ternary | NBR cover ternary | NBR_coverTernary | Full domain |
|  | NDVI cover ternary | NDVI_coverTernary |  |
|  | NIRv cover ternary | NIRV_coverTernary |  |
| Shape | NBR fractal dimension | NBR_FracDim | Disturbed pixels |
|  | NDVI fractal dimension | NDVI_FracDim |  |
|  | NIRv fractal dimension | NIRV_FracDim |  |
| Disturbance | NBR disturbance magnitude | NBR_mag | Disturbed pixels |
|  | NBR disturbance year of detection | NBR_yod |  |
|  | NBR disturbance signal-to-noise ratio | NBR_dsnr |  |
|  | NBR disturbance rate | NBR_rate |  |
|  | NDVI disturbance magnitude | NDVI_mag |  |
|  | NDVI disturbance year of detection | NDVI_yod |  |
|  | NDVI disturbance signal-to-noise ratio | NDVI_dsnr |  |
|  | NDVI disturbance rate | NDVI_rate |  |
|  | NIRv disturbance magnitude | NIRV_mag |  |
|  | NIRv disturbance year of detection | NIRV_yod |  |
|  | NIRv disturbance signal-to-noise ratio | NIRV_dsnr |  |
|  | NIRv disturbance rate | NIRV_rate |  |
| Texture | NBR contrast | NBR_distpx_tex_con | Disturbed pixels |
|  | NBR correlation | NBR_distpx_tex_cor |  |
|  | NBR entropy | NBR_distpx_tex_ent |  |
|  | NDVI contrast | NDVI_distpx_tex_con |  |
|  | NDVI correlation | NDVI_distpx_tex_cor |  |
|  | NDVI entropy | NDVI_distpx_tex_ent |  |
|  | NIRv contrast | NIRV_distpx_tex_con |  |
|  | NIRv correlation | NIRV_distpx_tex_cor |  |
|  | NIRv entropy | NIRV_distpx_tex_ent |  |

